# The spike-stabilizing D614G mutation interacts with S1/S2 cleavage site mutations to promote the infectious potential of SARS-CoV-2 variants

**DOI:** 10.1101/2022.05.20.492832

**Authors:** Stacy Gellenoncourt, Nell Saunders, Rémy Robinot, Lucas Auguste, Maaran Michael Rajah, Jerôme Kervevan, Raphaël Jeger-Madiot, Isabelle Staropoli, Cyril Planchais, Hugo Mouquet, Julian Buchrieser, Olivier Schwartz, Lisa A. Chakrabarti

## Abstract

SARS-CoV-2 remained genetically stable during the first three months of the pandemic, before acquiring a D614G spike mutation that rapidly spread worldwide, and then generating successive waves of viral variants with increasingly high transmissibility. We set out to evaluate possible epistatic interactions between the early occurring D614G mutation and the more recently emerged cleavage site mutations present in spike of the Alpha, Delta, and Omicron variants of concern. The P681H/R mutations at the S1/S2 cleavage site increased spike processing and fusogenicity but limited its incorporation into pseudoviruses. In addition, the higher cleavage rate led to higher shedding of the spike S1 subunit, resulting in a lower infectivity of the P681H/R-carrying pseudoviruses compared to those expressing the Wuhan wild-type spike. The D614G mutation increased spike expression at the cell surface and limited S1 shedding from pseudovirions. As a consequence, the D614G mutation preferentially increased the infectivity of P681H/R-carrying pseudoviruses. This enhancement was more marked in cells where the endosomal route predominated, suggesting that more stable spikes could better withstand the endosomal environment. Taken together, these findings suggest that the D614G mutation stabilized S1/S2 association and enabled the selection of mutations that increased S1/S2 cleavage, leading to the emergence of SARS-CoV-2 variants expressing highly fusogenic spikes.

**AUTHOR SUMMARY:** The successive emergence of SARS-CoV-2 variants is fueling the COVID pandemic, thus causing a major and persistent public health issue. The parameters involved in the emergence of variants with higher pathogenic potential remain incompletely understood. The first SARS-CoV-2 variant that spread worldwide in early 2020 carried a D614G mutation in the viral spike, making this protein more stable in its cleaved form at the surface of virions, and resulting in viral particles with higher infectious capacity. The Alpha and the Delta variants that spread in late 2020 and early 2021, respectively, proved increasingly transmissible and pathogenic when compared to the original SARS-CoV-2 strain. Interestingly, Alpha and Delta both carried mutations in a spike cleavage site that needs to be processed by cellular proteases prior to viral entry. The cleavage site mutations P681H/R made the Alpha and Delta spikes more efficient at viral fusion, by generating a higher fraction of cleaved spikes subunits S1 and S2. We show here that the early D614G mutation and the late P681H/R mutations act synergistically to increase the fusion capacity of SARS-CoV-2 variants. Specifically, viruses with increased spike cleavage due to P681H/R were even more dependent on the stabilizing effect of D614G mutation, which limited the shedding of cleaved S1 subunits from viral particles. These findings suggest that the worldwide spread of the D614G mutation was a prerequisite to the emergence of more pathogenic SARS-CoV-2 variants with highly fusogenic spikes.

## INTRODUCTION

The SARS-CoV-2 pandemic has remained a major public health issue during the past two years, due to the repeated emergence of viral variants endowed with increased transmissibility and/or immune escape capacity, leading to successive epidemic rebounds [1]. The SARS-CoV-2 virus is a beta-coronavirus characterized by a relative low mutation rate [2], but the sheer length of its RNA genome (30 kb) and its wide dissemination provide ample opportunities for the emergence and selection of viral mutations. SARS-CoV-2 remained genetically stable during the first few months of the pandemic, before the emergence in February-March 2020 of a variant characterized by a D614G mutation in the spike gene [3]. The spread of the D614G mutation proved remarkably rapid, as this mutation achieved worldwide dominance by June 2020 and has remained present in the vast majority of SARS-CoV-2 genomes sequenced so far [4]. The end of 2020 was marked by the independent emergence of divergent SARS-CoV-2 lineages: B.1.1.7 in the UK, B.1.351 in South Africa, and P1 in Brazil. These new lineages were associated with rapid epidemic rebounds, and have since then been renamed as the Alpha, Beta, and Gamma variants of concern (VOCs), respectively [5]. The emergence of the VOCs could be ascribed to an increase in transmissibility and in escape from innate immunity in the case of the Alpha variant [6–8], and to an efficient escape from preexisting neutralizing antibody responses in the case of the Beta and Gamma variants [9, 10]. Other emerging lineages characterized by a more localized spread, such as Kappa or Epsilon, were labelled as variants of interest (VOIs). February 2021 saw the emergence of the B.1.617 lineage in Maharashtra, India, with a particularly successful sublineage giving rise to the Delta VOC. The Delta variant proved even more transmissible and pathogenic than the Alpha variant, and ended up superseding all the preexisting variants by mid-2021 [11, 12]. In November 2021, a highly divergent VOC called Omicron emerged in South-Africa [13]. The transmissibility and immune escape capacity of Omicron proved superior to those of all the previous VOCs [14, 15], resulting into a worldwide replacement of Delta by Omicron by early 2022. Recent studies point to a lower pathogenicity of the Omicron variant [16], raising the possibility of a shift in SARS-CoV2 evolution towards viral attenuation.

Several key spike mutations have been shown to play a role in the increased transmissibility or immune escape capacity of the VOCs. The initial D614G mutation was shown to increase the infectivity of SARS-CoV-2 in several cell culture systems [3, 17]. This translated into an increased transmissibility and viral replication capacity in the hamster and ferret models of SARS-CoV-2 infection [18, 19]. Structurally, the D614G mutation abrogated an inter-protomer contact, which caused the spike receptor binding domain (RBD) to more readily adopt an up conformation, thus facilitating the interactions with the ACE2 receptor [20, 21]. Further structural studies showed that D614G also had a key role in stabilizing the non-covalent association of the S1 and S2 spike subunits once they were cleaved, through the ordering of a loop reinforcing intra-protomer interactions [17, 22]. Thus, the D614G mutation made the spike more stable but also more prone to interact with its receptor, accounting for the clear selective advantage conferred to the virus.

The N501Y mutation shared by the Alpha, Beta, and Gamma VOCs was shown to increase the affinity of the spike for the ACE2 receptor, and to be sufficient to increase SARS-CoV-2 infectivity and transmissibility in the hamster model [23]. Another group of spike mutations located close to the receptor binding motif, and including the E484K, K417N/T, and L452R substitutions, is involved in the escape of VOCs from neutralizing antibodies [1, 19]. A third group of VOC mutations are located in the vicinity of the S1/S2 cleavage site located at R685/S686, and may thus influence the processing of the spike. The original SARS-CoV-2 Wuhan strain is characterized by the presence of a polybasic motif 681-PRRAR-685 just N-terminal to the S1/S2 cleavage site, a feature unique among sarbecoviruses [24]. The polybasic motif is recognized by the furin protease during spike export to the surface of infected cells, resulting in a partial cleavage of the spike trimers incorporated into virion [25]. The cleavage at the S1/S2 site preactivates the spike, making SARS-CoV-2 virions less dependent on proteases expressed at the surface of target cells for their entry [26, 27] and more fit *in vivo* [28, 29]. Indeed, when the S1/S2 site is pre-cleaved, the spike bound to the ACE2 receptor requires only a single additional cleavage at the S2’ site to release the fusion peptide and transition to a fusogenic conformation [30]. This additional cleavage may be provided by the surface protease TMPRSS2, or by endosomal proteases such as cathepsin L if virions are endocytosed [4, 31, 32].

Notably, several of the SARS-CoV-2 variants carry a mutation at P681, just upstream of the polybasic S1/S2 cleavage site (Fig 1). The Alpha VOC and the Theta VOI carry a P681H change, which increase the local positive charge, and may thus facilitate furin cleavage [33]. Indeed, SARS-CoV-2 spikes carrying the P681H mutation were shown to be more cleaved and more fusogenic than their non-mutated counterparts in most [34–36], though not all studies [37]. Of note, the Alpha variant also carries a nearby T716I mutation that may further influence spike cleavage. The Delta variant and the Kappa VOI carry a P681R mutation, which further increases the basic nature of the S1/S2 cleavage motif. This mutation was shown to increase the degree of cleavage at S1/S2, resulting in a spike that is even more fusogenic than that of Alpha [38, 39]. The P681R mutation provides a clear competitive advantage to Delta in terms of infectivity in cell culture systems and in animal models of infection [38, 40]. More broadly, there has been a consistent trend for increased S1/S2 cleavage in the different VOCs and VOIs that emerged in the past two years, which may help account for their higher transmissibility and pathogenicity [35]. In contrast, the recently emerged Omicron variant shows a low degree of S1/S2 cleavage in virions released in culture, even though it carries the P681H mutation also found in the Alpha spike [14, 41]. The limited degree of precleavage reported for the Omicron spike, and its preference for the endosomal rather than the TMPRSS2 entry route, may contribute to the lower pathogenicity reported for this variant [16, 42].

**Fig 1.**
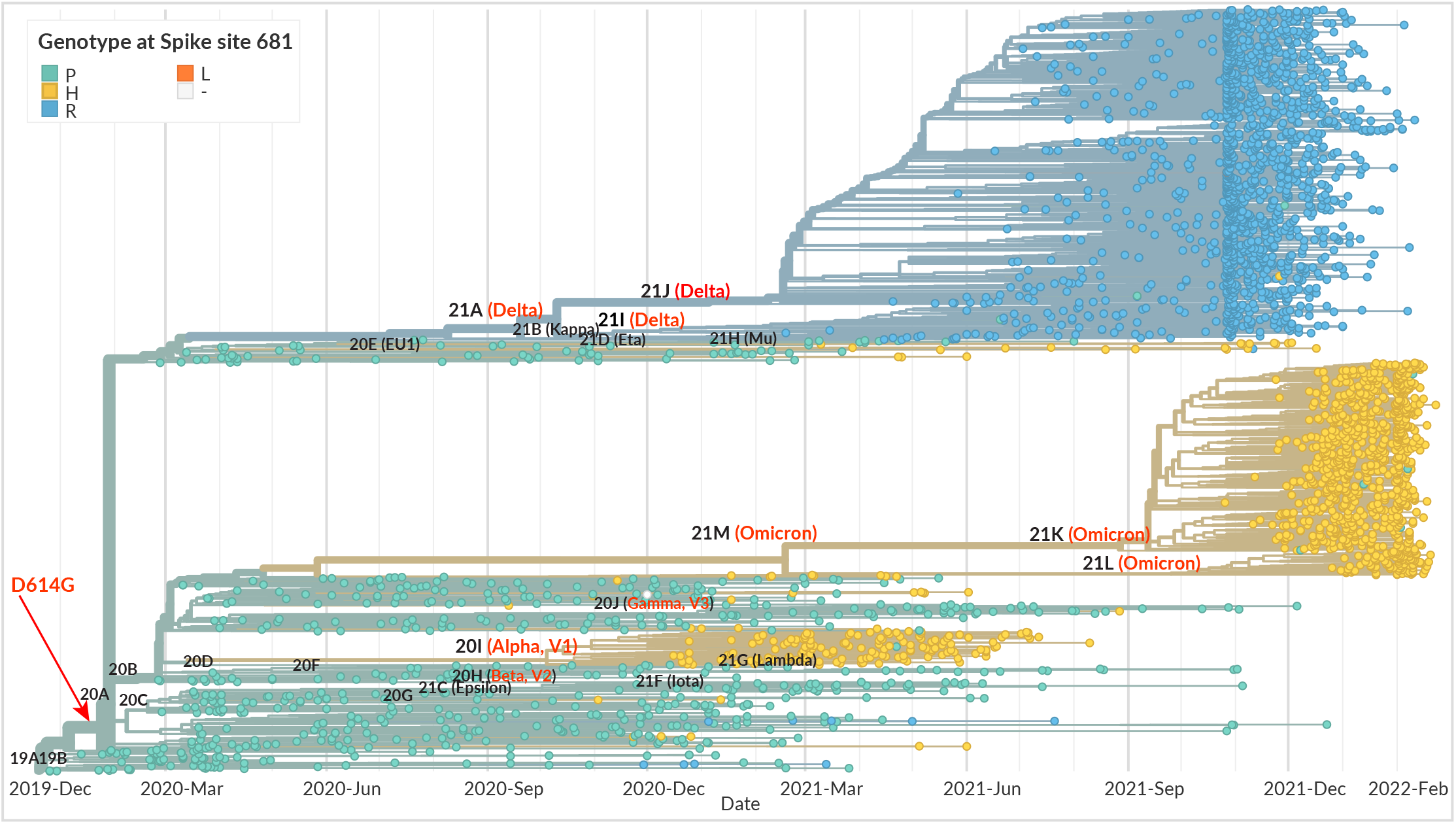
Phylogeny of SARS-CoV-2 variants, with spike genotype at position 681 highlighted. The phylogeny is based on 3,247 SARS-CoV-2 genomes sampled globally between Dec. 2019 and Feb. 2022, from the database curated by the GISAID Initiative (https://www.gisaid.org/). The phylogenetic tree was generated by the Nextstrain open source project (https://nextstrain.org/), with data points colorized according to spike genotype at position 681 (P681 turquoise; H681 yellow; R681 blue; L681 orange). The branches are labeled with SARS-CoV-2 clades according to the Nextrain nomenclature, with variant Greek names in parentheses. Variants of concern (VOCs) are highlighted in red. The estimated emergence of the D614G mutation is indicated by a red arrow.

Phylogenetic analyses suggest that each VOC arose independently, rather than by sequential evolution (Fig. 1). However, it remains intriguing that VOCs did not arise prior to the selective sweep that replaced the original Wuhan strain by the D614G-carrying variant. To address this issue, we assessed possible epistatic interactions between D614G and the dominant S1/S2 cleavage site mutations. Analyses of fusogenic capacity, spike processing, and infectivity of pseudotyped viruses did reveal interactions between these two types of genetic changes. Specifically, the stabilizing effect of the D614G mutation proved necessary to maintain the infectious capacity of virions carrying a highly fusogenic spike, pointing to the critical role of D614G in enabling the emergence of the VOCs.

## RESULTS

### Design of spike mutations

The cleavage site mutations present in the SARS-CoV-2 VOCs were introduced into the spike sequence of the reference Wuhan strain (Fig 2). Specifically, we constructed spike-expressing phCMV vectors carrying the punctual mutations P681H or T716I found in the Alpha variant, and P681R found in the Delta variant (Fig 2A). Of note, the P681H mutation is also present in the currently emerging Omicron variant. Controls included a Wuhan spike with a deletion of the furin cleavage site (FCS) equivalent to that found in the naturally occurring bat coronavirus RaTG13. This ΔFCS mutation corresponded to the deletion of aa 681-684 (PRRA), resulting in in a TNS/RSVA sequence at the S1/S2 junction. To evaluate interactions between the early occurring D614G mutation and the more recently emerged cleavage site mutations, the Wuhan (WT) and cleavage site mutant spikes were constructed in two versions, with or without the D614G mutation (Fig 2B). As references, we also generated spikes carrying the full complement of mutations present in the Alpha and Delta variants.

**Fig 2.**
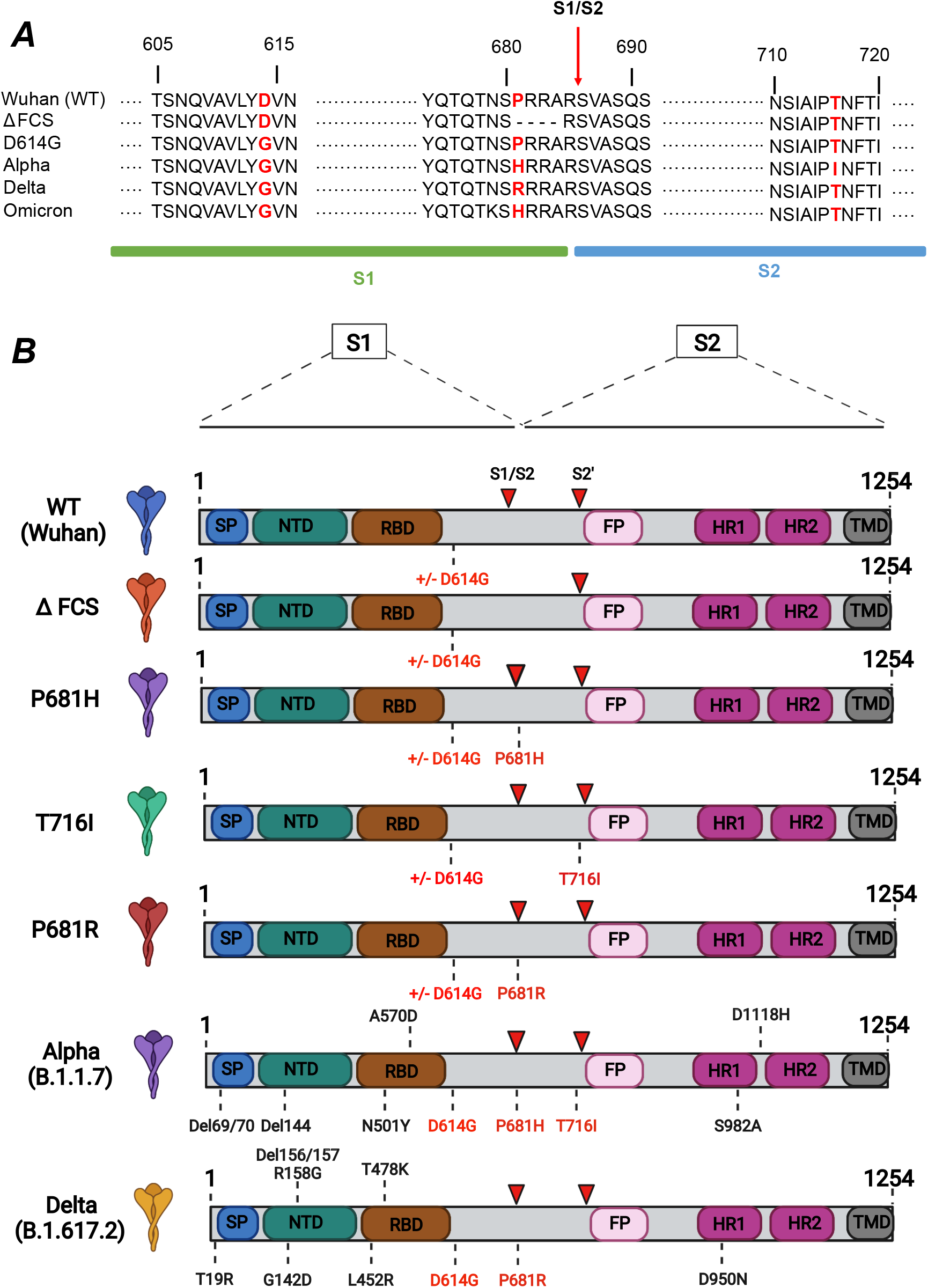
Spike mutations analyzed in the present study. (A) Sequences surrounding the spike mutations studied. The S1/S2 spike cleavage site is indicated by a vertical arrow. The mutations studied are highlighted in red. The ΔFCS mutation corresponds to the deletion of the PRRA sequence at the S1/S2 cleavage site. (B) Schematic of the spike expression vectors used in the present study. The mutations ΔFCS, P681H, P681R, and T716I were inserted into the Wuhan WT spike backbone truncated at residue 1254, in the presence or absence of the D614G mutation. The spikes containing the full complement of mutations present in the Alpha and Delta VOCs were used as controls. SP: signal peptide; NTD: N-terminal domain; RBD: receptor binding domain; FP: fusion peptide; HR: heptad repeat; TMD: transmembrane domain. Red arrowheads indicate the S1/S2 and S2’ cleavage sites.

### The D614G mutation and the deletion of the furin cleavage site both increase spike expression at the cell surface

Spike-expressing vectors were initially transfected in HEK 293Tn cells (hereafter HEK cells) to verify that the spike proteins were properly expressed at the cell surface. Transfected cells were labelled with the human monoclonal antibody mAb 129 known to cross-react with the S1 subunit of the SARS-CoV-2 variants included in the study [36]. All of the vector tested resulted in efficient expression of the spike, with over 70% of HEK cells labeled with the mAb 129 antibody (Fig 3A-B and S1A Fig). Analysis of the mean fluorescence intensity (MFI) of transfected HEK cells showed that the ΔFCS spike deleted of the FCS was expressed to higher levels (Fig 3C), possibly because the S1 subunit could not be shed from the cell surface. The difference in MFI between the WT and ΔFCS spikes proved significant only for the versions of the spikes expressing G614. To systematically evaluate the effect of the D614G mutation on spike surface expression, we computed the ratios of MFI for spike pairs carrying G614 or D614 (Fig 3E). This analysis showed that mean MFI ratios were above 1, suggesting that G614 increased spike surface expression. The MFI ratio was significantly higher for WT spikes than P681R spikes, suggesting that the D614 effect was modulated by spike cleavage. Similar trends were observed when analyzing the G614/D614 ratios of the percentage of spike-expressing cells (Fig 3D), though the difference between WT and P681R did not reach significance. Overall, this analysis showed that both the FCS deletion and the D614G mutation promoted spike expression at the cell surface.

**Fig 3.**
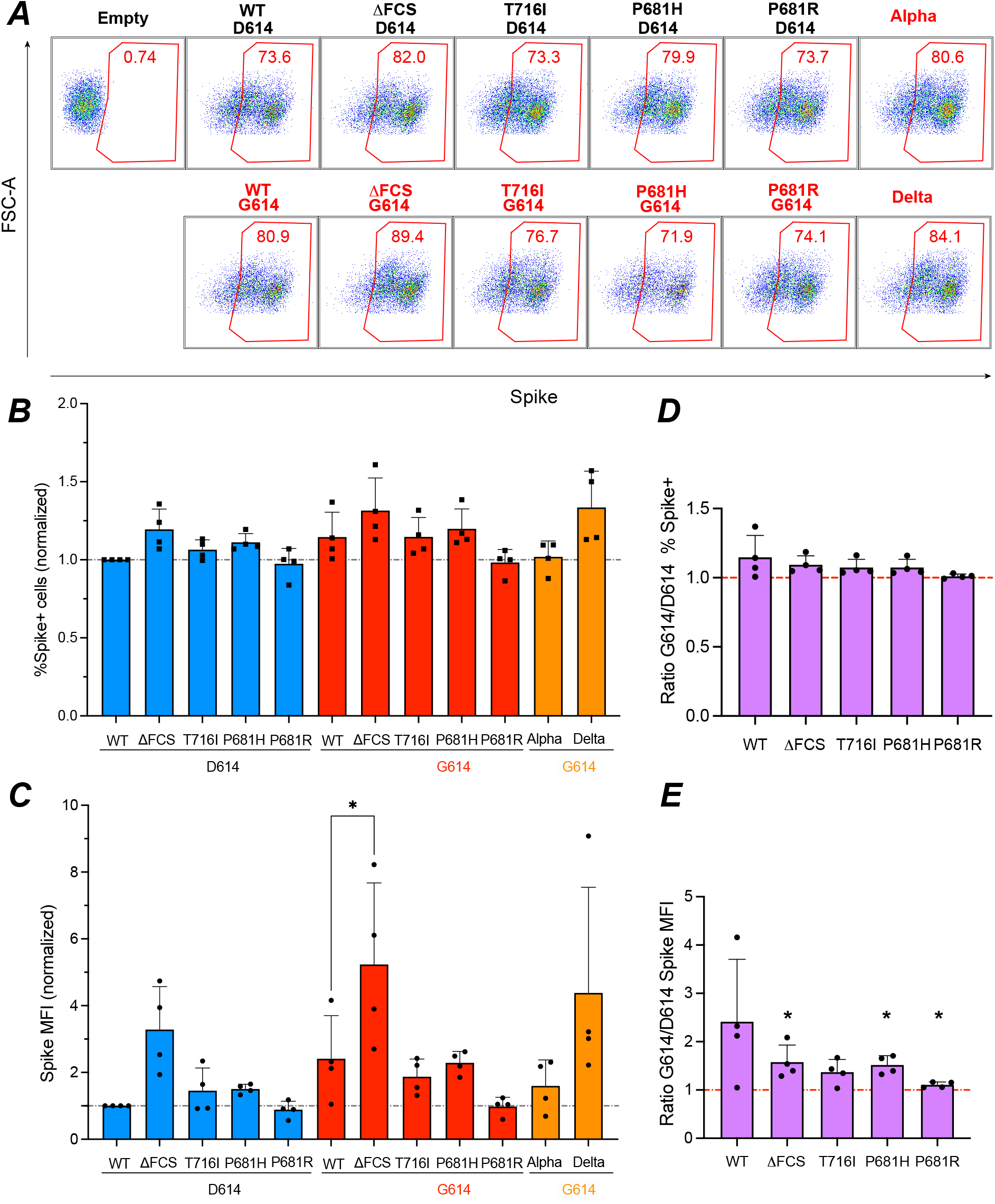
Increased spike surface expression upon deletion of the furin cleavage site. (A) Representative examples of spike surface expression. HEK 293T cells were transfected with spike expression vectors and analyzed by flow cytometry after surface staining with the spike-specific human mAb 129. The percentage of spike+ cells in live cells is reported. (B, C) The percentage (B) and mean fluorescence intensity (MFI) of spike+ cells (C) normalized to WT is reported. Statistics were measured by one-way ANOVA, with the Holm-Sidak’s correction for multiple comparisons. Mutants in the D614 backbone were compared to the WT D614 spike, while mutants in the G614 backbone were compared to the WT G614 spike. (D, E) The ratio of spike surface expression measured in the G614 backbone to that in the D614 backbone is reported, using either the percentage (D) or the MFI of spike+ cells (E) for measurement. Statistics evaluating whether each ratio is different from 1 were measured by one sample t-tests. (B-E) Means +/− SD are shown for n=4 independent experiments. * p<0.05.

### The P681H/R mutations increase SARS-CoV-2 spike fusogenicity

The fusogenic capacity of the different spikes was evaluated in a GFP-split system where spike-transfected HEK cells expressing the truncated reporter protein GFP1-10 were mixed with Vero-E6 cells expressing the complementary reporter protein GFP11. Upon cellular fusion, reconstitution of a functional GFP protein due to GFP1-10/GFP11 interaction resulted in fluorescent emission at 488 nm (Fig 4A). GFP+ syncytia were visualized on an Opera imaging system and the GFP+ surface normalized to the number of nuclei was quantified by automated image analysis.

**Fig 4.**
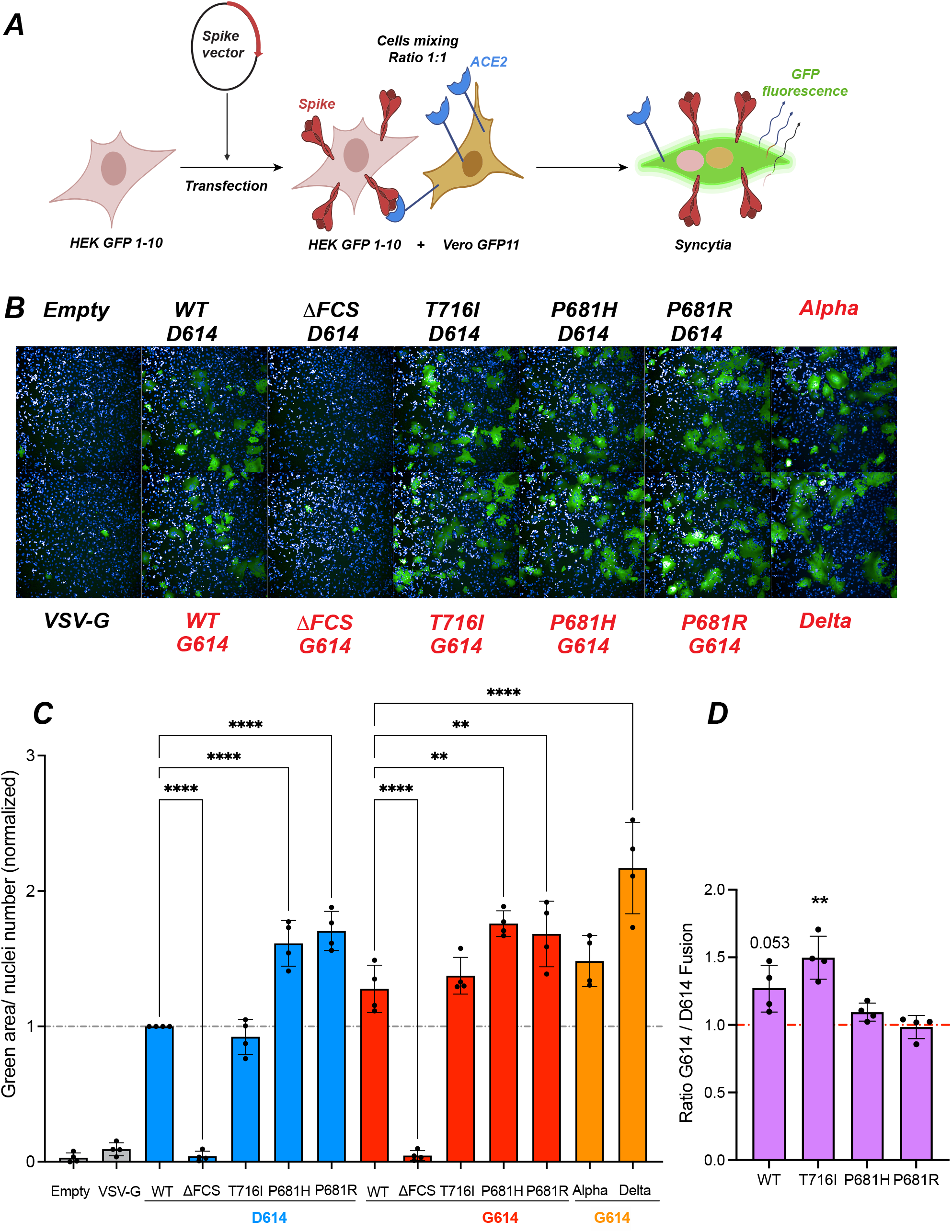
The P681H/R mutations increase spike fusogenicity. (A) Principle of the cell-cell fusion assay. HEK GFP1-10 are transfected with spike vectors and then mixed with Vero GFP11 at a 1:1 ratio. Spike-ACE2 interactions lead to cell-cell fusion. Upon syncytia formation, complementation between the GFP1-10 and GFP11 fragments generate functional GFP, resulting in a fluorescent emission. (B) Representative images of a fusion assay monitored at 18h post-spike transfection. Cell nuclei were stained with DAPI (blue) while syncytia were detected by GFP fluorescent emission (green). The spike vectors used as reported, with names colorized in red for those containing the D614G mutation. (C) Quantification of cell-cell fusion induced upon spike expression. The green fluorescent area divided by the number of nuclei is reported, with normalization to the WT spike condition. Statistics were measured by one-way ANOVA, with the Holm-Sidak’s correction for multiple comparisons. Mutants in the D614 backbone were compared to the WT D614 spike, while mutants in the G614 backbone were compared to the WT G614 spike. (D) The ratio of cell-cell fusion measured for spikes in the G614 backbone to that in the D614 backbone is reported. Statistics evaluating whether each ratio is different from 1 were measured by one sample t-tests. (C,D) Means +/− SD are shown for n=4 independent experiments. Each dot represents the mean of technical triplicate measurements. * p<0.05; ** p<0.01; *** p<0.001; **** p<0.0001.

The ΔFCS spike did not induce detectable fusion (Fig 4B), compatible with the notion that cleavage at the FSC is required for SARS-CoV-2 spike-dependent fusion at the cell surface. In contrast, all of the other spikes induced the formation of GFP+ syncytia, with the size of syncytia appearing larger for spikes carrying the P681H/R mutations (Fig 4B). Quantitative image analysis showed that the WT and T716I spikes had equivalent fusogenic capacity (Fig 4C). In contrast, the P681H and P681R spikes induced significantly more fusion than the WT spike (P<0.0001 in presence of D614; P<0.01 in the presence of G614), consistent with previous findings by us and others [35, 36]. The Alpha and Delta spikes also proved strongly fusogenic, compatible with the presence of the P681H and P681R mutations, respectively.

Comparison of fusion achieved in presence and absence of the D614G mutation was measured by fusion ratios (Fig 4D). This analysis showed that the D614G mutation increased the fusogenicity of the WT Wuhan and T716I spikes, with ratios ≥1, possibly due to increased spike expression at the cell surface. In contrast, the D614G effect was weak or absent for the already highly fusogenic spikes carrying P681H/R mutations.

### Cleavage site mutations limit spike incorporation into pseudoviruses

Pseudotyped viruses (PV) were produced by co-transfecting HEK cells with a SARS-CoV-2 spike expression vector, a GFP-lentivector backbone, and HIV-derived packaging plasmids. PV particles concentrated by ultracentrifugation were analyzed for spike incorporation and processing by Western blotting. Dual labeling was obtained with a polyclonal anti-S1 antibody (green fluorescence) and a monoclonal anti-S2 (red fluorescence), so that the uncleaved spike precursor S0 containing both the S1 and S2 subunits appeared as yellow (Fig 5A, top panel). PV protein extracts were normalized according to their content in HIV p24 capsid protein (Fig 5A, bottom panel).

**Fig 5.**
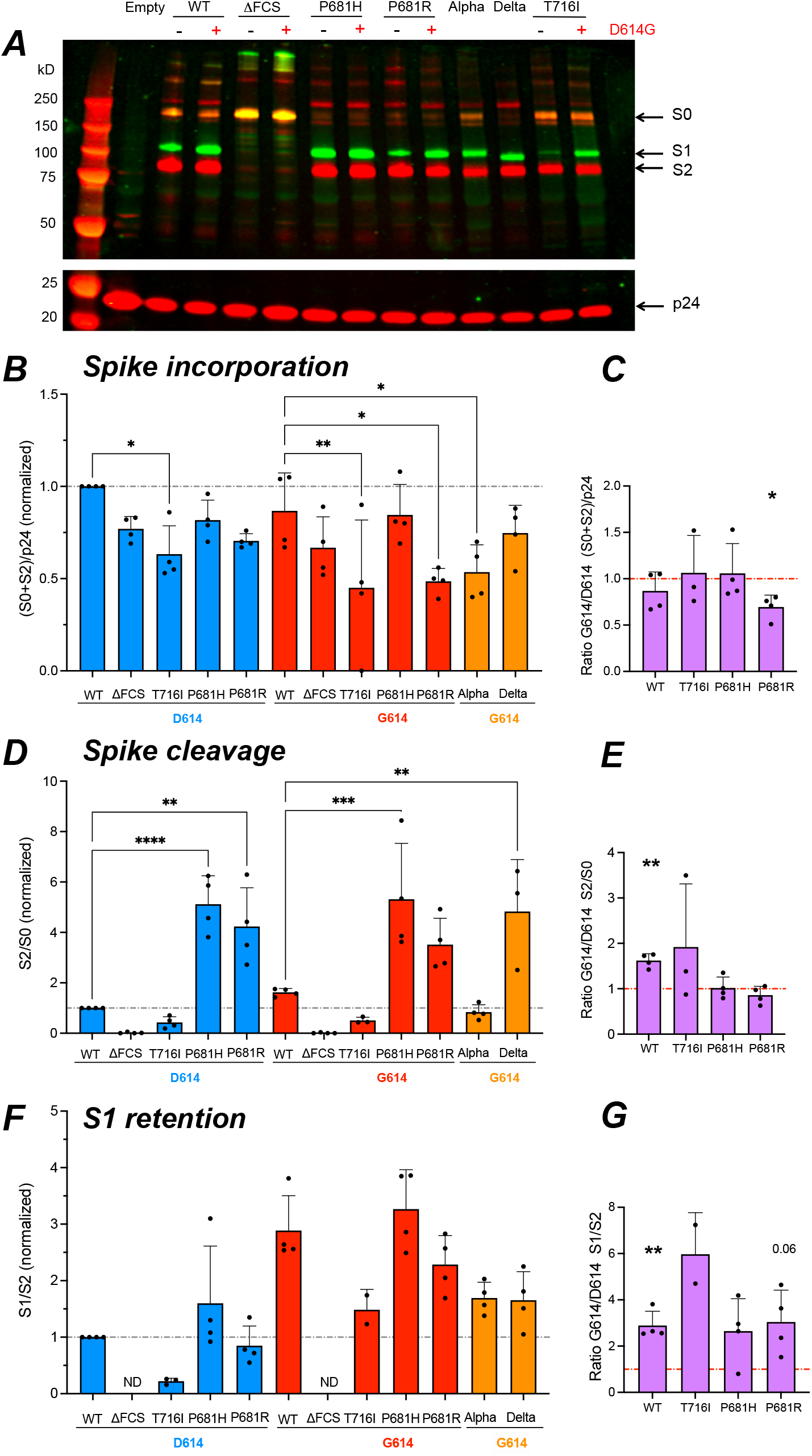
The D614G mutation promotes S1 subunit retention in virions. The different forms of spike present into pseudotyped viruses (PV) were analyzed by Western blotting on concentrated PV particles. (A) Representative Western blot showing the S1 (green) and S2 (red) spike subunits in the top panel, and the p24 Gag protein (red) used for normalization in the bottom panel. The equivalent of 300 ng p24 was loaded in each lane. Molecular weight markers were loaded in the first lane, with the expected marker size in kD indicated on the left. The uncleaved spike precursor S0 is visible in yellow due to the superposition of green and red fluorescence. (B) Quantitation of total spike incorporated (S0 + S2), reported to p24 content, and normalized to WT. The fluorescence intensity of each band was quantified with the Image Studio Lite software. (C) Ratio of total spike incorporated measured in the G614 backbone to that in the D614 backbone. (D) Quantitation of total cleaved spike, measured by the S2/S0 ratio, and normalized to WT. (E) Ratio of total cleaved spike measured in the G614 backbone to that in the D614 backbone. (F) Quantitation of S1 subunit retention in PV particles, measured by the S1/S2 ratio, and normalized to WT. ND: not detectable. (G) Ratio of S1 retention measured in the G614 backbone to that in the D614 backbone. (B, D, F) Statistics were measured by one-way ANOVA, with the Holm-Sidak’s correction for multiple comparisons. Mutants in the D614 backbone were compared to the WT D614 spike, while mutants in the G614 backbone were compared to the WT G614 spike. (C, E, G) Statistics evaluating whether each ratio was different from 1 were measured by one sample t-tests. (B-G) Means +/− SD are shown for n≥3 independent experiments, except for T716I S1 retention, where n=2. * p<0.05; ** p<0.01; *** p<0.001; **** p<0.0001.

Particle pseudotyped with the ΔFCS spike showed a highly predominant S0 band, as expected for a spike devoid of an S1/S2 cleavage site. In contrast, S1 and S2 subunits were detected in PV expressing a spike with an FCS. To evaluate the total amount of spike incorporated by the different PV, we measured the summed intensities of the S2 and S0 bands (in the red channel), reported to the amount of p24 capsid protein (Fig 5B). Of note, the (S0+S2)/p24 parameter takes into account all the spikes moieties incorporated into the PV, irrespective of their cleavage status or of the possible shedding of the S1 subunit. Quantitation of this parameter showed that all of the cleavage site mutations tended to decrease total spike incorporation, with an effect that was more marked for the T716I mutant. The Alpha PV also showed low spike incorporation compared to the WT G614 PV. Based on a G614/D614 ratio analysis (Fig 5C), the D614G mutation had minimal effect on total spike incorporation, except for a possible decrease in incorporation of the P681R.

### The D614G mutation promotes spike cleavage but limits S1 shedding

We then analyzed the ratio of S2 subunit to S0 precursor, in order to evaluate the extent of spike cleavage in PV particles. This analysis showed that the P681H/R mutations strongly promoted spike cleavage (Fig 5D), consistent with the literature [34, 35, 38, 39]. The Delta PV also harbored almost fully cleaved spikes, while this was not apparent for the Alpha PV, suggesting that other mutations such as T716I counterbalanced the pro-cleavage effect of P681H. The G614/D614 ratio analysis showed that the presence of the G614 mutation increased spike cleavage in the WT and T716I PV, but had no discernable effect on the already highly cleaved spikes of the P681H and P681R PV (Fig 5E).

The relative amount of S1 subunit retained on the different PV was then measured by quantifying the S1/S2 ratio (Fig 5F). The T716I mutation was associated to a very low S1/S2 ratio relative to WT, while the P681H/R mutations did not have a marked effect on the S1/S2 ratio. Thus, the T716I mutation induced marked S1 shedding, while the addition of a basic amino acid in the FCS appeared neutral on shedding, even though it increased spike cleavage. Presence of the D614G mutation promoted S1 retention in all the PV tested, as shown by values above 2 in the G614/D614 ratio analysis relative to the S1/S2 parameter (Fig 5G). In particular, D614G efficiently corrected the deleterious effect of T716I on S1 shedding. Taken together, the D614G mutation promoted spike cleavage but limited S1 shedding, enabling the production of viral particles with a high content of cleaved spike.

A Western blot analysis was also carried out on lysates of HEK cells used to produce the PV particles (S2 Fig). The parameters that measured spike content ((S0+S2)/actin), spike cleavage (S2/S0), and S1 shedding (S1/S2) showed patterns similar to those observed in PV, indicating that the studied spike mutations already exerted their effects at the level of producer cells. One exception was the lower cleavage of the spike carrying both the P681R and D614G mutations in producer cells as compared to viral particles (S2C Fig). This observation suggested that spike carrying P681R could be further cleaved after viral budding.

### The D614G mutation preferentially increases the infectivity of P681H/R-carrying pseudoviruses

To evaluate the impact of spike mutations on viral infectivity, we measured the extent of single-cycle infection induced by PV on HEK cells expressing the ACE2 receptor, in presence or absence of the TMPRSS2 coreceptor (S1B Fig). At 48h, the infection was evaluated by the percentage of target cells expressing the GFP reporter gene transferred by the pseudotyped lentivector (representative examples in Fig 6A and S3 Fig). This analysis showed that, in absence of the D614G change, the infectivity of PV carrying the cleavage site mutations T716 and P681H/R was low as compared to WT (Fig 6B). In contrast, the ΔFCS spike conferred high infectivity in HEK-ACE2 cells, indicating that an FCS was not required for viral entry in these cells. A dose response analysis confirmed these findings at different viral input doses (S4 Fig). Infectivity was overall higher in HEK-ACE2-TMPRSS2 compared to HEK-ACE2 cells (S4A Fig). Differences in infectivity between the PV were less marked than in HEK-ACE2 cells (Fig 6B-C), suggesting that an additional entry route enabled by the TMPRSS2 protease could attenuate mutant-induced entry defects.

**Fig 6.**
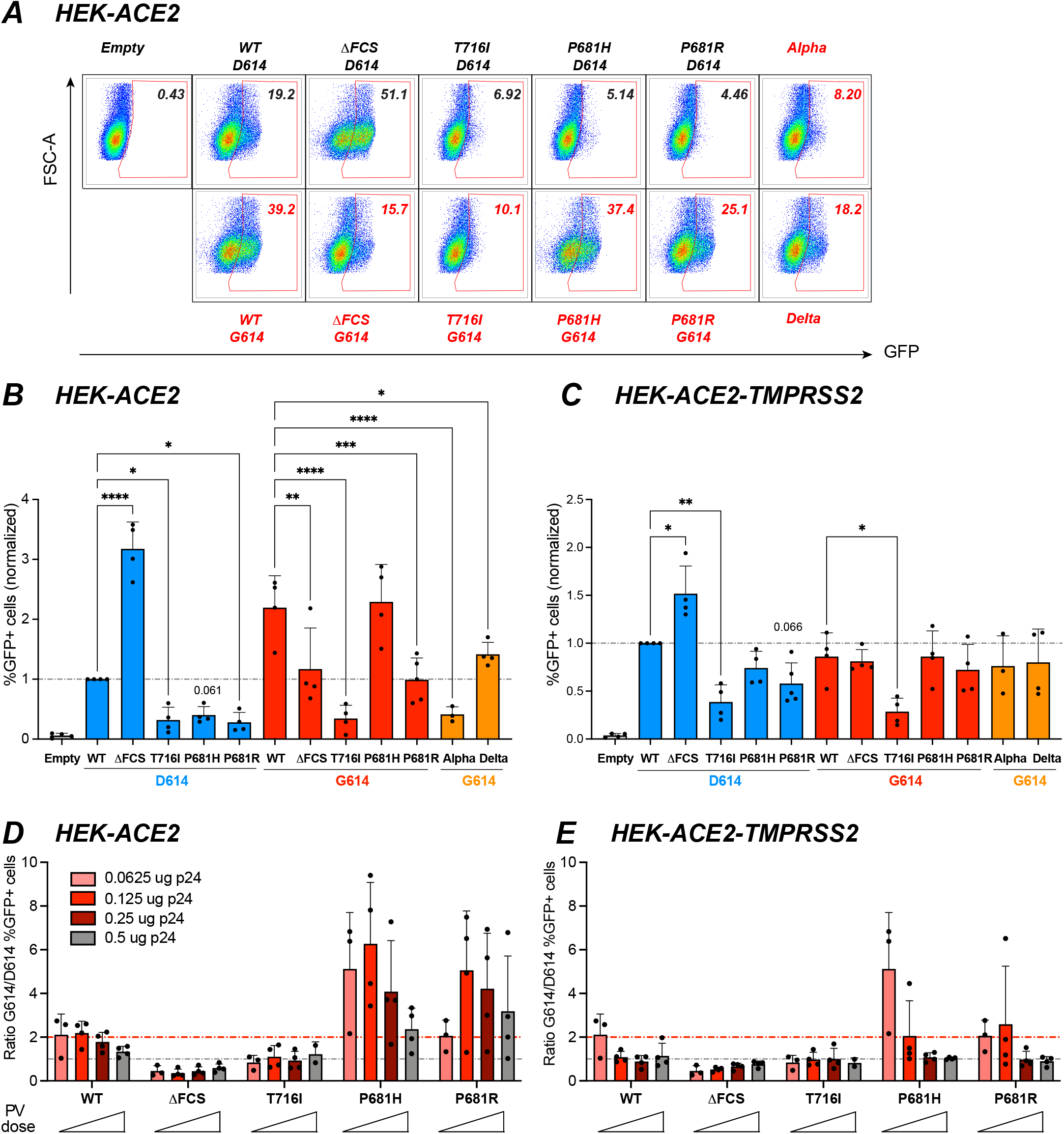
The D614G mutation preferentially increases the infectivity of P681H/R-carrying pseudoviruses. The infectivity of GFP-lentivector pseudotyped with the different spikes was analyzed in HEK-ACE2 cells induced or not for TMPRSS2 expression. (A) Representative infection experiment in HEK-ACE2 cells. Cells were inoculated with 0.25 μg of p24 equivalent. The infectivity of pseudoviruses (PV) is measured by the percentage of GFP+ cells reported in the top right corner of each plot. (B, C) Quantitation of PV infectivity at the intermediate PV dose corresponding to 0.125 μg of p24 equivalent. The percentage of GFP+ cells obtained in HEK-ACE2 cells (B) and HEK-ACE2-TMPRSS2 cells (C) normalized to WT is reported. Statistics were measured by one-way ANOVA, with the Holm-Sidak’s correction for multiple comparisons. Mutants in the D614 backbone were compared to the WT D614 spike, while mutants in the G614 backbone were compared to the WT G614 spike. Means +/− SD are shown for n≥3 independent experiments. (D, E) The ratio of PV infectivity in HEK-ACE2 (D) and in HEK-ACE2-TMPRSS2 (E) measured for spikes in the G614 backbone to that in the D614 backbone is reported. The ratio was computed for increasing doses of PV ranging from 0.0625 to 0.5 μg pf p24 equivalent. The preferential effect of the D614G mutation on the infectivity of PV carrying the P681H/R mutation is evidenced by ratios above 2 (red line: ratio=2; grey line: ratio=1). Means +/− SD are shown for n≥3 independent experiments, except for the T716I mutant at the highest dose where n=2.

Analysis of the G614/D614 infectivity ratios showed that the D614G mutation increased infectivity of the WT, P681H, and P681R PV in HEK-ACE2, while this was not the case for the T716I and ΔFCS PV (Fig 6D). The enhancement of viral infectivity tended to be more marked at lower PV input doses, presumably because of saturation of the readout and a degree of cytotoxicity observed at the highest input dose (S4B Fig). Interestingly, the effect of D614G was more marked for the PV carrying the basic P681R/H mutations than for the WT, as shown by ratios ≥2 (Fig 6D, red line), suggesting that D614G may have played a compensatory role that was important in the emergence of the P681H/R-carrying variants. The Alpha PV which retained relatively low infectivity in HEK-ACE2 cells (Fig 6B), suggesting that mutations other than D614G and P681H also impacted viral entry, with a possible negative role of T716I. The effect of D614G was less apparent in HEK-ACE2-TMPRSS2 than in HEK-ACE2 cells, but remained more marked for the P681R/H than the WT PV (Fig 6C,E), confirming that D614G preferentially increased the infectivity of PV carrying the P681H/R basic mutations.

### The D614G mutation increases infectivity in different cellular contexts

To test the generality of these findings, we measured the infectivity of PV in the osteosarcoma cell lines U2OS-ACE2 +/− TMPRSS2 and in the lung adenocarcinoma cell line Calu-3. As PV expressing a GFP reporter gene gave a low detection signal in these cell lines, we switched to a PV system where the backbone lentivector expressed luciferase, resulting in a higher signal to noise ratio (Fig 7). The pattern of infectivity observed in U2OS-ACE2 was similar to that observed in HEK-ACE2, with a relative defect of the T716I, P681H and P681R PV compared to WT, and a highly efficient entry of the ΔFCS PV (Fig 7A). The D614G mutation markedly increased PV infectivity in U2OS-ACE2 cells (Fig 7B), with a partial restoration of infectivity for the P681H/R mutants. The Alpha PV showed an infectivity equivalent to that of WT-G614, while that of the Delta PV proved significantly higher (Fig 7A).

**Fig 7.**
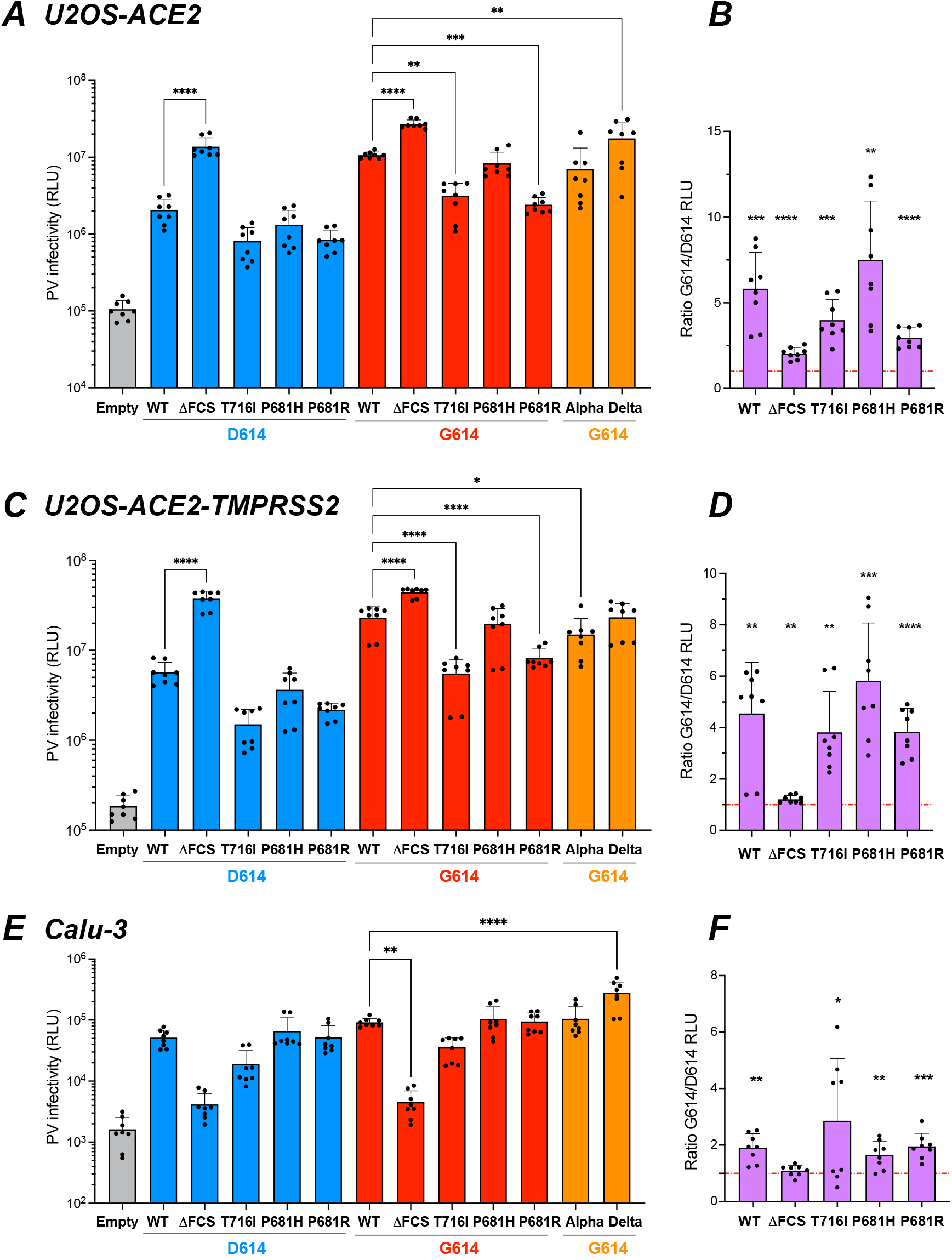
The D614G mutation increases infectivity in different cellular contexts. The infectivity of Luc-lentivector pseudotyped with the different spikes was analyzed in different cells lines. Results are expressed in Relative Luciferase Units (RLU). (A-D) Luciferase activity in U2OS-ACE2 cells (A, B) and U2OS-ACE2-TMPRSS2 cells (C,D) infected with 0.125 μg pf p24 equivalent. (E-F) Luciferase activity in Calu-3 cells infected with 1 μg pf p24 equivalent. (A, C, E) Statistics were measured by one-way ANOVA, with the Holm-Sidak’s correction for multiple comparisons. Mutants in the D614 backbone were compared to the WT D614 spike, while mutants in the G614 backbone were compared to the WT G614 spike. (B, D, F) The ratio of PV infectivity in U2OS-ACE2 (B), U2OS-ACE2-TMPRSS2 (D), and Calu-3 cells (F) measured for spikes in the G614 backbone to that in the D614 backbone is reported. Statistics evaluating whether each ratio is different from 1 were measured by one sample t-tests. (A-F) Means +/− SD are shown for n=4 independent experiments, with 2 technical replicates per experiment. * p<0.05; ** p<0.01; *** p<0.001; **** p<0.0001.

Analyses in U2OS-ACE2-TMPRSS2 showed that the presence of TMPRSS2 led to an overall increase in PV infectivity (Fig 7C). The relative differences in PV infectivity showed a pattern similar to that observed in absence of TMPRSS2, except that the Delta PV showed an infectivity comparable rather than higher than WT. Presence of the D614G mutation led to a full restoration of infectivity for the P681H PV, and to a partial restoration for the P681R PV (Fig 7C, D).

In the Calu-3 cell line, which expresses ACE2 and TMPRSS2 endogenously [32], PV entry was lower than in cell lines transduced with the ACE2 and TMPRSS2 receptor (Fig 7E). The ΔFCS PV showed drastically decreased infectivity in Calu-3, indicating that a functional FCS was required for efficient entry in this cell line. The T761I PV still showed a decreased infectivity as compared to WT, while the P681H/R PV were as infectious as WT in the Calu-3 setting. The Alpha PV also showed an infectivity comparable to WT-G614, while infectivity of the Delta PV was significantly increased (Fig 7E).

The G614/D614 infectivity ratios were above 1 for all the PV tested (Fig 7F), indicating that the D614G mutation increased infectivity in Calu-3 cells, though this increase appeared moderate compared to that seen in other cell lines. Concerning the P681R/H mutants, the infectivity ratios were in the 1.5-2.5 range in Calu-3, while they were in the 3-7 range HEK-ACE2 and U2OS-ACE2 +/− TMPRSS2, suggesting that the compensatory effect of the D614G mutation depended on the cellular context.

### Limited effect of the D614G mutation on entry pathway usage

SARS-CoV-2 is known to use different entry pathways depending on the availability of the proteases involved in spike processing. Specifically, expression of TMPRSS2 promotes entry at the cell surface, while in the absence of this protease, viral entry can take place in endosomes after spike processing via cathepsins B and L [4, 31, 32]. As we noted a variable compensatory effect of the D614G mutation depending on the target cell type, we asked whether D614G influenced entry pathway usage. To this goal, target cells were pretreated with the serine protease inhibitor Camostat, which blocks TMPRSS2 activity, and/or with the cysteine protease inhibitor E64D, which blocks the activity of the endosomal cathepsins B and L [31]. Pretreated cells were then infected and cultivated in the presence of the inhibitors for two days before evaluating PV infectivity by luciferase activity.

Viral entry in HEK-ACE2 cells was inhibited at 80% by E64D treatment for most PV tested, while the blocking effect of Camostat was minimal (20%), suggesting that the endosomal entry route predominated in this cell line (Fig 8A and S5A Fig). The combination of E64D and Camostat resulted in an inhibition equivalent to that seen with E64D alone, reinforcing the notion of a predominant endosomal entry. Entry inhibition did not vary depending on the presence or absence of the D614G mutation, suggesting that spike stability was not the main parameter determining the viral entry route. The Delta PV was characterized by a lower susceptibility to entry blocking by the two protease inhibitors, possibly due to the high fusogenicity of the Delta spike. Another exception was the T716I PV carrying the D614G mutation, which showed only 60%inhibition by E64D treatment. However, this resistance was only relative, as this PV retained an overall low infectivity in HEK-ACE2 compared to the other PV carrying D614G (S5A Fig).

**Fig 8.**
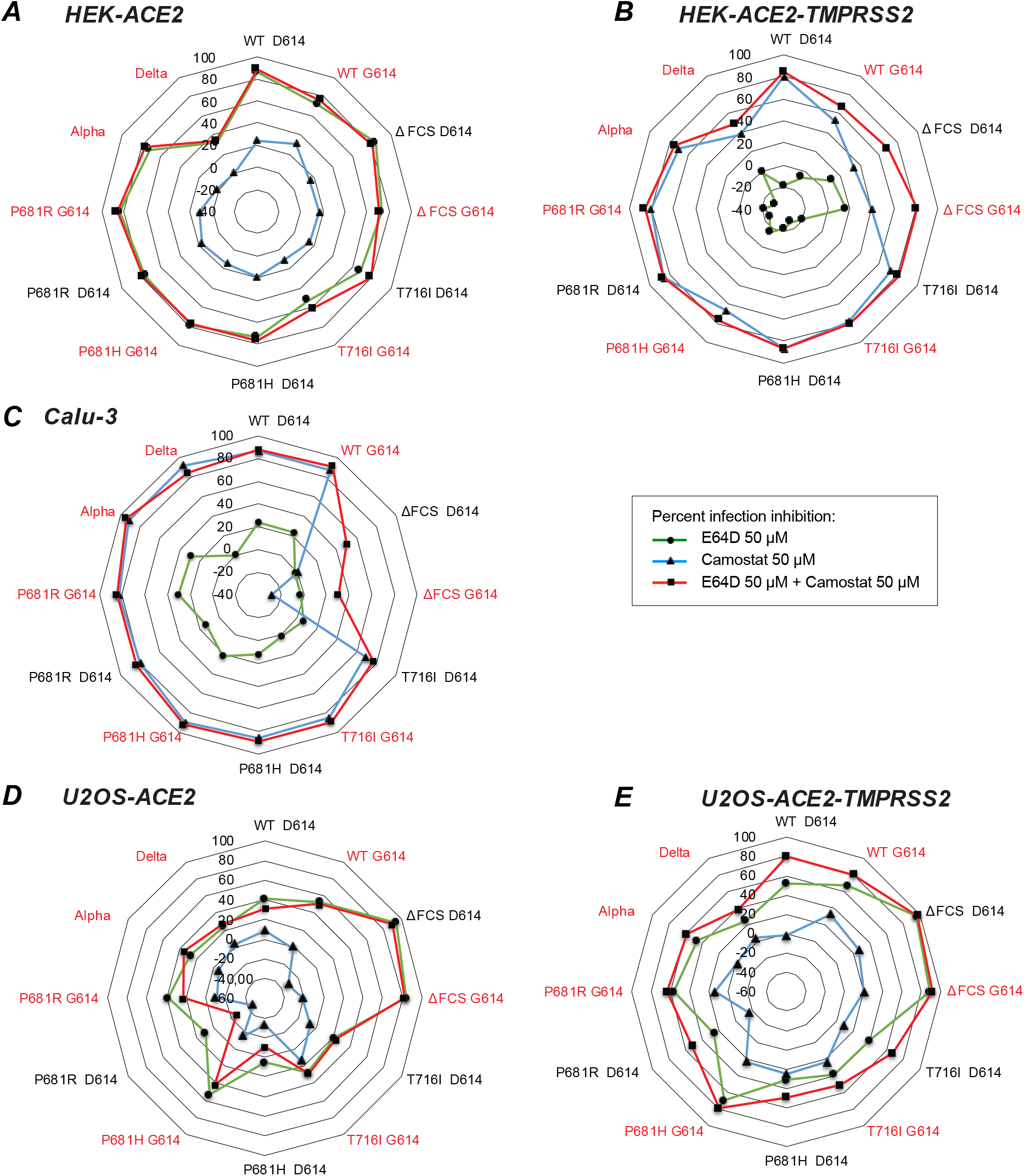
Partial inhibition of infectivity by combined TMPRSS2 and cathepsin inhibitors in U2OS cells. The TMPRSS2 inhibitor Camostat and the cathepsin inhibitor E64D evaluated for their capacity to block pseudovirus (PV) infectivity, alone or in combination. The percentage of infectivity inhibition was measured by the mean decrease in luciferase activity (1-(RLU treated/RLU untreated)) x 100 for n=3 independent experiments, with 2 technical replicates per experiment. The percentage of infectivity inhibition is reported in a radar plot for Camostat (blue line with triangles), E64D (green line with circles), or the combination of the two inhibitors (red line with squares) in the following cell lines: HEK-ACE2 (A), HEK-ACE2-TMPRSS2 (B), Calu-3 (C), U2OS-ACE2 (D), and U2OS-ACE2-TMPRSS2 (E). The names of PV carrying the D614G mutation are reported in red.

Analysis of entry inhibition in HEK-ACE2-TMPRSS2 showed an inverted pattern as compared to that seen in HEK-ACE2, with about 80% inhibition by Camostat, and a negligible effect of E64D (Fig 8B). Thus, the expression of TMPRSS2 was sufficient to switch entry pathways, with the surface route becoming predominant. The Delta PV proved again less susceptible to entry inhibition. The other PV that showed low (40%) entry inhibition by Camostat was ΔFCS, consistent with the limited capacity of an uncleaved spike to promote surface entry. The ΔFCS PV were the only ones to show a degree of inhibition by E64D in HEK-ACE2-TMPRSS2, suggesting that a limited access to the endosomal route remained available in the presence of TMPRSS2. The presence of the D614G mutation had little or no effect on susceptibility to protease inhibitors, suggesting that the main determinants of entry pathway usage lied with the host cell rather than with the viral particle.

Evaluation of entry inhibition in Calu-3 cells yielded a pattern that was overall similar to that seen in HEK-ACE2-TMPRSS2, compatible with a predominant use of the surface entry route in both cell lines (Fig 8C and S5C Fig). Camostat proved highly efficient at blocking viral entry in Calu-3, with inhibition percentages above 80% for most PV, and no further increase in inhibition when adding E64D to Camostat. The exception was again the ΔFCS PV, which were not inhibited by Camostat. However, entry of the ΔFCS PV was only marginally higher than that seen for the control PV devoid of spike (Empty, S5C Fig), confirming that a spike devoid of the FCS enabled only minimal entry in Calu-3 cells.

### Partial inhibition of infectivity by combined TMPRSS2 and cathepsin inhibitors in U2OS cells

The entry inhibition pattern seen in U2OS-ACE2 cells proved intermediate, with neither Camostat nor E64D achieving efficient blocking of viral entry in this cell line (Fig 8D-E and S5D-E Fig). The inhibition induced by E64D was in the 20-40% range, except for ΔFCS PV, which remained cathepsin-dependent for viral entry. We noted that PV carrying G614 tended to be more susceptible to E64D inhibition than their D614 counterpart, suggesting that PV with stabilized spikes may still partially use the endosomal entry pathway in the U2OS cellular context. Camostat did not inhibit viral entry in U2OS-ACE2 and, intriguingly, showed also a limited effect (about 20% inhibition) in U2OS-ACE2-TMPRSS2 (Fig 8E). In contrast, the inhibitory effect of E64D was maintained and even at time increased (20-60% range) in the presence of TMPRSS2. When considering absolute infectivity values, addition of TMPRSS2 did increase viral entry in U2OS-ACE2 cells (S5E Fig). However, the expression of TMPRSS2 was not sufficient to entirely redirect viral entry to a Camostat-dependent pathway, in contrast to findings obtained in HEK-ACE2-TMPRSS2 cells. The partial entry inhibition observed with combined Camostat plus E64D treatment raised the possibility of an additional entry route that would not depend on TMPRSS2 nor cathepsins.

## DISCUSSION

This study demonstrates that spikes carrying the S1/S2 cleavage site mutations characteristic of SARS-CoV-2 variants require the additional D614G mutation to maintain an efficient viral entry. The acquisition of the D614G mutation was already viewed as a key event that enabled the worldwide dissemination of the original SARS-CoV-2 strain [3, 4]. We report here that D614G is even more important for the fitness of virions carrying the P681R/H mutations, in particular when the endosomal entry pathway predominates. Indeed, the benefit conferred by D614G on infectivity was 1.5 to 3 times higher for pseudoviruses carrying the P681R/H mutations than for those with the original P681 residue in HEK-ACE2 cells. Based on these findings, we propose that acquisition of D614G played an essential role in the emergence of the highly transmissible and pathogenic Alpha and Delta variants.

We had previously shown that P681H was the key mutation involved in conferring a higher fusogenicity to the Alpha spike [36]. We report in the present study that the P681R mutation found in the Delta spike confers an even higher fusogenic capacity, consistent with the recent literature [38, 40, 43]. In a D614 context, these highly fusogenic spikes were characterized by equivalent or slightly lower expression than the Wuhan spike at the surface of producer cells, and by decreased incorporation into pseudotyped virions. This may be due to differences in spike trafficking, as premature intracellular fusion events triggered by the P681H/R spikes could perturb protein maturation pathways. Introduction of the D614G mutation slightly increased the surface expression of the P681H spike, but not that of the P681R spike, and did not restore incorporation of these spikes into virions. Thus, the D614G change must have acted primarily after the virion release stage. The P681H/R spikes incorporated into virions showed a clear increase in the degree of cleavage at the S1/S2 site, as expected [17, 39, 40]. Introduction of the D614G mutation into the P681H/R spikes did not significantly change their degree of cleavage, as measured by the S2/S0 ratio. The key difference proved to be in the degree of S1 subunit retention on virions, with an S1/S2 ratio increased by a factor 2 to 3 in the presence of the D614G mutation. Of note, increased S1 retention was also seen when D614G was inserted into the original Wuhan spike, as reported here and in previous studies [17, 22]. However, due to the higher cleavage rate achieved by the P681H/R spikes, the proportion of S1 subunits retained in presence of the D614G was higher, accounting for the preferential effect of the D614G change on highly fusogenic spikes.

The infectivity enhancement conferred by the D614G mutation proved higher in the absence of TMPRSS2 overexpression. This was observed in the HEK-ACE2 as well as the U2OS-ACE2 cellular contexts. In addition, the infectivity enhancement remained moderate in Calu-3 cells, which endogenously express TMPRSS2 and enable viral entry via the surface rather than the endosomal route [32]. A likely explanation for these findings is that S1 subunit stability is more critical when using the endosomal rather than the TMPRSS2-dependent surface entry pathway. SARS-CoV-2 entry via the endosomal route was shown to take longer than via the surface route, and to require endosomal acidification [31, 32]. A more stable spike is likely to better withstand a long entry process and/or an acidifying environment. Ultrastructural studies suggest that the release of S1 subunits facilitates the transition of the metastable spike trimer to a post-fusion conformation, and that these changes may at times happen prematurely, before attachment of S1 to its receptor ACE2 [4]. The presence of post-fusion trimers lacking S1 has for instance been documented at the surface of virions by cryo-electron microscopy [44], and it is possible that this phenomenon is further triggered within endosomes. The D614G mutation may limit these premature conformational changes, as this mutation was shown to stabilize the non-covalent association of the S1 and S2 subunits within a spike protomer [22]. The stabilizing effect of the D614G mutation may thus be particularly relevant in the case of pre-cleaved spikes with a high propensity for fusion, and in cellular contexts that favor the endosomal entry route.

A spike devoid of the polybasic S1/S2 cleavage site (ΔFCS) proved defective in the cell-cell fusion assay. In contrast, pseudotypes carrying this spike were highly infectious in the HEK-ACE2 and U2OS-ACE2 cellular contexts, consistent with previous reports [25, 28, 45]. These observations highlight a disconnect between fusogenic capacity measured at the cell surface and infectivity, likely due to differences in the subcellular locations involved. Indeed, the ΔFCS spike was highly sensitive to E64D inhibition but only minimally sensitive to camostat, and thus mediated entry almost exclusively by the endosomal route. This was further confirmed by the low infectivity of ΔFCS pseudotypes in Calu-3 cells where the surface route predominates, as previously reported [27]. The ΔFCS spike is deleted of the PRRA sequence, but still retains an arginine at 685, and may thus be cleaved at this site (or at neighboring sites) by cathepsins, accounting for its capacity to mediate entry through endosomes. Of note, presence of the D614G mutation did not restore the fusogenic capacity of the ΔFCS spike, and induced only limited changes in its infectivity pattern. Indeed, D614G tended to decrease, rather than increase, the infectivity of ΔFCS pseudotypes in HEK-ACE2 cells, slightly increased infectivity in U2OS-ACE2 cells, and had no effect on infectivity in Calu-3 cells. These observations further support the notion that D614G acts through stabilization of S1/S2 association, as D614G loses its effects in situations where S1 and S2 are covalently associated.

The sensitivity of pseudotyped viruses to entry inhibitors did not appear dependent on the D614G mutation. Thus, D614G modulated the magnitude of infection, but not the proportion of infection that took place through the surface or the endosomal entry routes. Rather, the choice of entry route varied according to the cell line studied, suggesting that it mainly depended on host cell characteristics. As expected, the expression of TMPRSS2 redirected most of entry through the surface route, in HEK-ACE2-TMPRSS2, as well as Calu-3 cells. The findings in U2OS-ACE2 cells were more intriguing, as neither camostat nor E64D could fully block pseudotyped virus entry in this cell line (with the exception of ΔFCS). Further, expression of TMPRSS2 in these cells did not fully redirect viral entry to a camostat-sensitive pathway. These observations raise the possibility of an additional entry pathway that would be insensitive to both camostat and E64D, and would thus not depend on TMPRSS2 nor on cathepsins. This putative pathway would still require a polybasic S1/S2 cleavage site, as it did not play a role in the entry of the ΔFCS pseudotyped virus. One may note that alternate receptors have been recently proposed for SARS-CoV-2 [46], which may provide a basis for alternate entry routes.

The T716I mutation found in the Alpha spike had an overall deleterious effect on pseudotyped virus infectivity, which was associated with decreased spike incorporation into virions. In addition, the T716I mutation induced slightly less spike cleavage, but markedly lower retention of cleaved S1 subunits. Taken together, the T716I change resulted in a low amount of functional spike carrying the S1 subunit on virions, accounting for a low infectivity. These observations are compatible with structural studies showing that the T716I mutation has a locally destabilizing effect, by abrogating an intra-spike protomer hydrogen bond [47]. Combination of D614G with the T716I mutation did have a stabilizing effect, as S1 retention improved. However, the infectivity of the D614G/T716I pseudotypes remained lower than that of other mutants in all the cell lines tested. The presence of T716I change may help explain why Alpha pseudotypes retain a low infectivity in HEK-ACE2 cells. On the other hand, Alpha pseudotypes are as infectious as WT G614 pseudotypes in U2OS-ACE2 and Calu-3 cells, implying that the deleterious effect of T716I is compensated by other mutations present in the Alpha spike. Reasons for the conservation of a destabilizing mutation such as T716I remain to be fully elucidated, but may be related to the capacity of the Alpha spike trimer to more frequently switch to an RBD up conformation competent for ACE2 binding [47]. Overall, structural studies point to the interplay of several stabilizing and destabilizing mutations that control the degree of exposure of the RBD.

Compensatory mutations that increase entry of the Alpha spike act through several mechanisms. A two residues deletion found in the N-terminal domain of the Alpha spike, ΔH69V70, was reported to increase total spike incorporation into virions [48]. In contrast, the D614G changes acts primarily through the stabilization of the S1/S2 non-covalent association, as discussed above. The N501Y change found in the RBD increases affinity of the spike for ACE2 by a factor up to 5, which promotes viral entry [23, 47].The P681H mutation increased spike fusogenicity, which should have an impact on viral entry. However, P681H by itself did not increase infectivity in our assays, consistent with a previous report [34]. Possible reasons include the instability of the cleaved S1 subunit, which may not be fully compensated by the D614G change, and a trend for lower incorporation of the P681H spike into viral particles. While the P681H change does not directly increase infectivity of the Alpha variant, it may contribute its pathogenic potential though the facilitation of cell-cell fusion. A pathogenic role for syncytia in pulmonary alveoli has been proposed [43, 49], which may help account for increased pathogenicity of the Alpha variant compared to that of the original Wuhan strain [50]. The increased in fusogenicity associated to P681R mutation is even more marked, which may contribute to the superior pathogenicity of the Delta variant [12]. The recently emerged variant Omicron expresses a spike with reduced fusogenic capacity [14, 41] and has shown signs of decreased pathogenicity [16], further supporting and association between these two parameters.

It remains intriguing that the Omicron variant retains the P681H change in its spike, in spite of its low fusogenic capacity and a reported low degree of S1/S2 cleavage [41]. Omicron carries 33 non-synonymous spikes mutation compared to the Wuhan strain, an unprecedented number of changes that induce a remodeling of the NTD and RBD surfaces, and result in immune evasion from most classes of monoclonal antibodies [51]. These changes also impact the viral entry step, as Omicron infectivity is hampered in conditions of low ACE2 expression [52], a property that may explain the low infectivity of Omicron for lung alveolar tissue [42]. Structurally, the regions that control RBD flipping appear more structured in the Omicron trimer, which may slow down the transition of the RBD to the up position, and thus make ACE2 binding a rate limiting step in viral entry [52]. Residues close to the FCS also appear more structured, which may limit access of the furin enzyme, and account for the low cleavage rate. Thus, low fusogenicity of Omicron appears to result from combinatorial changes that restrict several steps of the entry process, but accommodate a wide array of immune escape mutations.

The SARS-CoV-2 virus is demonstrating remarkable adaptability in the face of worldwide vaccination efforts and mounting natural immunity. Due to the plasticity of the spike, emerging variants could adopt distinct evolutionary strategies, based on different degrees of trade-off between transmissibility, pathogenicity, and immune escape. It remains striking, however, that all the SARS-CoV-2 variants relied on the D614G mutation for S1 subunit stabilization. We report that the dependency on the D614G mutation was more marked for highly fusogenic spikes, implying a key role of the D614G mutation in the emergence of highly pathogenic SARS-CoV-2 variants.

## MATERIALS AND METHODS

### Plasmids

All the spike mutations were inserted into a codon-optimized version of the Wuhan-Hu-1 SARS-CoV-2 spike (GenBank: QHD43416.1) cloned into a phCMV backbone (GenBank: AJ318514). Mutations were introduced into the phCMV-Spike plasmids using the mutagenic primers and the Q5 site directed mutagenesis kit (NEB). In addition, a truncation of 19 amino acids was introduced into the cytoplasmic tail of each spike studied (deletion at amino acids 1255 to 1273 in the Wuhan-Hu-1 spike), to promote spike incorporation into lentiviral particles, as described previously [53, 54]. All the mutant plasmids were sequenced prior to use. Primers used for mutagenesis and sequencing are reported in S1 Table. The plasmid pQCXIP-empty was used as a negative control for spike expression [55]. Plasmids used to produce GFP-lentiviruses were the lentivector backbone pCDH-EF1α-GFP (System Biosciences), the packaging plasmid psPAXII (Addgene), and the pRev plasmid (a gift from P. Charneau). For the production of luciferase-lentivirus, the plasmids used were pHAGE-CMV-Luc2-IRES-ZsGreen (#NR-52516), pHDM-Hgpm2 (#NR-52517), pHDM-tat1b (#NR-52518), pRC-CMV-rev1b (#NR-52519), all generated in J. Bloom’s laboratory [56] and obtained from BEI Resources (kit NR-53816).

### Cell lines

HEK 293Tn (purchased from SBI Biosciences) and Calu-3 (ATCC #HTB55) cell lines were maintained in Dulbecco’s modified Eagle medium (DMEM) and DMEM-F12, respectively, supplemented with 10% fetal bovine serum and 100 μg/mL penicillin/streptomycin (DMEMc). Cell lines transduced with ACE2 and/or TMPRSS2 have been previously described [55]. HEK 293T-hACE2-TMPRSS2 cells (called herein HEK-ACE2-TMPRSS2) were induced for TPMRSS2 expression by addition of doxycycline (0.5 μg/mL, Sigma) and were maintained in DMEMc with blasticidin (10 μg/mL, InvivoGen) and puromycin (1 μg/mL, Alfa Aesar). U2OS cells expressing hACE2 and GFP1-10 or hACE2, TMPRSS2, and GFP1-10 were maintained in DMEMc supplemented with blasticidin (10 μg/mL, InvivoGen), puromycin (1 μg/mL, Alfa Aesar) and hygromycin B (0.2 mg/mL). HEK 293T and Vero-E6 cells expressing GFP1-10 and GFP11 were maintained in DMEMc supplemented with 1 μg/mL and 4 μg/mL of puromycin, respectively. All the cell lines were cultured at 37°C under a 5% CO2 atmosphere, and were routinely screened for mycoplasma.

### Transfection of HEK 293Tn cells with spike vectors

Transfection was performed with Lipofectamine 2000 (ThermoFisher, #11668019), using 125 ng of phCMV-Spike plasmid diluted in OptiMem medium in a final volume of 25 μL. DNA was incubated with 0.9 μL Lipofectamine 2000 diluted in 25 μL OptiMem medium was added to the DNA, and the mixture was incubated for 20 minutes, before being added onto 0.1 million of HEK 293Tn cells seeded into a 96-well plate. Transfection efficiency was assessed 18h post-transfection by monitoring spike surface expression by flow cytometry.

### Production of spike-pseudotyped lentivectors

GFP lentiviral particles pseudotyped with the SARS-CoV-2 spike were prepared by transfection of HEK 293Tn cells using the CaCl2 method. The lentiviral vector pCDH-EF1a-GFP, the packaging plasmid psPAXII, the spike expression vector phCMV-Spike and the pRev plasmid were mixed at a 2:2:1:1 ratio, with a total DNA amount of 252 μg used per 175 cm^2^ flask. The pQCXIP-empty plasmid was used to generate a spike-negative control. At 48h post-transfection, supernatants were collected and spun for 5 min at 270 g for 5 min at 4°C to remove cell debris. Lentiviral particles were then concentrated by ultracentrifugation at 23,000 g for 1h30 at 4°C onto a 20% sucrose cushion. The viral particle pellet was resuspended in PBS and frozen in aliquots at −80°C until use.

Luciferase lentiviral particles were produced according to the same protocol, but with different plasmid ratios: the lentiviral backbone pHAGE-CMV-Luc2-IRES-ZsGreen, the packaging plasmid pHDM-Hgpm2, the Tat and Rev plasmids pHDM-tat1b and pRC-CMV-rev1b, and the spike plasmid phCMV-Spike were used at ratio 4.4:1:1:1:1.5 to transfect a 175 cm^2^ flask. The pseudotyped lentiviral particles were normalized according to their capsid content, by measuring the Gag p24 antigen concentration with the Alliance HIV-1 p24 Antigen ELISA kit (Perkin Elmer).

### Infection with spike-pseudotyped lentivectors

The day before infection with spike-pseudotyped GFP-lentivectors, 100,000 HEK-ACE2-TMPRSS2 cells were plated in 96-well plates, and TMPRSS2 was induced when needed by the addition of doxycycline. HEK-ACE2 +/− TMPRSS2 were infected with serial 2x dilutions of GFP-lentivectors, ranging from 0.0625 μg to 0.5 μg of p24 Gag content in a final volume of 100 μL. Cells were cultivated in presence of lentivectors, and infection was quantified at 2 days post-infection (p.i) by measuring the percentage of GFP+ cells by flow cytometry.

For infection with spike-pseudotyped luciferase lentivectors, the cells were plated in flat-bottom 96-well plates the day before infection at the following concentrations: 20,000 U2OS GFP1-10 ACE2 +/− TMPRSS2, or 50,000 HEK 293T ACE2 +/− TMPRSS2, or 50,000 Calu-3 cells. HEK and U2OS were infected with 0.125 μg of p24 Gag equivalent, while Calu-3 were infected with 1 μg of p24 Gag equivalent, all in a final volume of 100 μL. Two days p.i, medium was removed and replaced by 30 μl DMEM without red phenol. The same volume of luciferase Bright-Glo substrate (Promega) was added before acquisition of luminescence on a Wallac Victor2 1420 Multilabel counter (Perkin Elmer system, v3.00.0.53). Infectivity was expressed in relative luciferase units (RLU). For infections experiments in the presence of protease inhibitors, cells were pretreated for 2 hours before infection with 100 μM of E64-D (Enzo Life Sciences, #BML-PI107-0001), or 100 μM of Camostat (CliniSciences, #HY-13512), or both, in a final volume of 50 μL. For infection, 50 μL of medium containing 0.065 μg of p24 Gag equivalent was added onto HEK ACE2 +/− TMPRSS2 and U2OS ACE2 +/− TMPRSS2, while 0.5 μg of p24 Gag equivalent in 50 μl added onto Calu-3 cells, resulting in a 2x dilution of the protease inhibitors. Infection was monitored at 2 days p.i by measuring luciferase activity, as described above. The percentage of infectivity inhibition was computed as follows: (1 - (RLU treated / RLU untreated)) x100.

### Western blotting

To prepare protein extracts, 2 millions of cells were lysed in buffer containing 150 mM NaCl, 50 mM Tris HCl (pH8), 1% Triton, 5mM EDTA, with 1x protease inhibitor cocktail (Roche, #11873580001) for 30 min on ice. For lentiviral particle extracts, an equivalent of 300 ng of Gag p24 was lysed in ELISA lysis buffer containing 1% Triton (Alliance HIV-1 p24 Antigen ELISA kit, Perkin Elmer) and protease inhibitors for 30 min on ice. Samples were heated and reduced in 1x final DTT buffer (Thermofisher, #NP0004) at 90°C for 10 min before being run on a 4-12% acrylamide denaturing gel (ThermoFisher, #NP0323 NuPAGE). Proteins were then transferred onto a nitrocellulose membrane (ThermoFisher, #IB23001). The membrane was blocked with 5% dried milk in PBS Tween 0.1 %, before incubation with three primary antibodies (S1, S2, and p24 Gag or actin) for 1h at room temperature (RT), followed by 3 washes in PBS Tween 0.1 %, and incubation with secondary antibodies for 30 min at RT. After 3 more washes, the fluorescent signal was revealed on a Li-Cor Odyssey CLx 9140 imaging system. Images were quantified with the Image Studio Lite (v5.2.5) software, using a mode with automated background subtraction. Primary antibodies consisted in two anti-spike antibodies (rabbit anti-S1 Genetex #GTX135356, 1:1000 and mouse anti-S2 Genetex #GTX632604, 1:1000), and one normalization antibody: mouse anti-p24 Gag (R&D Systems # MAB7360; 1:1000) or mouse anti-actin (Cell Signaling #8H10D10, 1:2000). Anti-rabbit and anti-mouse IgG secondary antibodies, conjugated to DyLight-800 (Bethyl Laboratories, #A80-304D8) and DyLight-680 (ThermoFisher, #SA5-35521), respectively, were used at a 1:5000 dilution each.

### Cell-cell fusion assay

For the cell-cell fusion assay, 15,000 GFP1-10-expressing HEK 293T were transfected with phCMV-Spike vectors, washed with DMEM supplemented with 10% FCS without antibiotics (DMEMna), and then mixed at a 1:1 ratio with GFP-11-expressing Vero-E6 cells. Briefly, 10 ng of spike plasmid and 90 ng of empty vector were mixed in 25 μl OptiMem medium. Then, 0.3 μL Lipofectamine 2000 diluted in 25 μl OptiMem medium was added to the DNA, and the mixture was incubated for 30 min before being added onto HEK GFP 1-10 cells. After 30 min, transfected cells were washed in DMEMna and spun at 500 g for 5 min before being mixed at a 1:1 ratio with Vero GFP11 cells in DMEMna. Cells were stained with Hoechst. Cells were cultivated for 18 hours in transparent bottom plates (μClear, #655090). Fusion measurement was assessed on an Opera Phenix High-Content Screening System (PerkinElmer), by automatically measuring number the GFP area normalized to the total number of nuclei.

### Flow cytometry

To quantify cells infected by GFP lentivectors, cells were harvested, washed in PBS, and stained with the viability dye Aqua (Invitrogen) at a 1:800 dilution in PBS for 30 min at 4°C. After two washes in PBS, cells were fixed with 2% paraformaldehyde (PFA) (ThermoFisher, #43368-9M) and acquired on an Attune NxT flow cytometer. To quantify spike expression in transfected HEK 293Tn cells, cells were harvested as above and stained with the pan-SARS-CoV-2 S1 human mAb129 [36] at 1 μg/mL and the viability dye Aqua for 30 min at 4°C. After two PBS washes, cells were stained with a Goat anti-human AF647 antibody (ThermoFisher, #A21445) at 1:500 for 30 min at 4°C. Samples were fixed with 2% PFA before acquisition on the NxT flow cytometer. Data were analyzed with the FlowJo software v10.7.1 (Becton Dickinson).

To evaluate ACE2 and TMPRSS2 expression in the target cell lines used, cells were surface-labelled with the goat anti-ACE2 antibody (R&D #AF933) at 5 mg/mL for 30 min at 4°C. After washing, cells were incubated with the secondary antibody donkey anti-goat-AF647 (ThermoFisher #A21447) at a 1:500 dilution. Cells were then permeabilized with the Fix/Perm kit (BD Biosciences #554714) according to manufacturer recommendations. Cells were intracellularly stained with a rabbit anti-TMPRSS2 antibody (Atlas antibodies, HPA035787) at 0.3 mg/mL. After washing, the secondary staining was done with a donkey anti-rabbit-AF555 antibody (Fisher #16229260) at a 1:500 dilution. Cells were then fixed in 2% PFA and analyzed by flow cytometry as described above.

### Statistical analysis

All statistical analyses were carried out with the GraphPad Prism software (v9). The tests used and significance thresholds are indicated into the legend of each figure. Mutants in the D614 backbone were compared to the WT D614 spike, while mutants in the G614 backbone were compared to the WT G614 spike. Statistics were computed for all spikes studied by one-way ANOVA with a Holm-Sidak correction for multiple comparisons. For G614/D614 ratio analyses, we evaluated by one sample t-test whether each ratio significantly differed from 1. All the graphs were generated in GraphPad Prism, except for the spider plots, which were made using Microsoft Excel (v16.16.27).

## ACKNOWLEDGEMENTS

We thank Nathalie Aulner and the UTechS Photonic BioImaging (UPBI) core facility of Institut Pasteur, a member of the France BioImaging network, for training on the Opera imaging system. We thank Jacob Seeler for help with gel image acquisition and all the members of the Virus and Immunity Unit for reagents and helpful discussions. The following reagent was obtained through BEI Resources, NIAID, NIH: “SARS-Related Coronavirus 2, Wuhan-Hu-1 Spike-Pseudotyped Lentiviral Kit V2, NR-53816.”

## FUNDING

This work was supported by: the Urgence COVID-19 Fundraising Campaign of Institut Pasteur (COROCHIP, PFR4, and PFR7 projects), and Agence Nationale de Recherche sur le SIDA et les Maladies Infectieuses Emergentes (ANRS-MIE; project 19206) (L.A.C.); the Pasteur Institute, the European Health Emergency Preparedness and Response Authority (HERA), Urgence COVID-19 Fundraising Campaign of Institut Pasteur, Fondation pour la Recherche Médicale (FRM), ANRS-MIE, the Vaccine Research Institute (ANR-10-LABX-77), Labex IBEID (ANR-10-LABX-62-IBEID), ANR/FRM Flash Covid PROTEO-SARS-CoV-2, ANR Coronamito, and IDISCOVR (O.S). S.G. is the recipient of a MESR/Ecole Doctorale BioSPC, Université Paris Cité fellowship, R.R. of a Sidaction fellowship, and J.K. of an ANRS fellowship.

## AUTHOR CONTRIBUTIONS

**Conceptualization**: Stacy Gellenoncourt, Olivier Schwartz, Lisa A. Chakrabarti

**Funding acquisition**: Olivier Schwartz, Lisa A. Chakrabarti

**Investigation**: Stacy Gellenoncourt, Nell Saunders, Rémy Robinot, Lucas Auguste, Maaran Rajah, Jérôme Kervevan, Raphaël Jeger-Madiot, Isabelle Staropoli

**Methodology**: Julian Buchrieser

**Resources**: Cyril Planchais, Hugo Mouquet

**Supervision**: Olivier Schwartz, Lisa A. Chakrabarti

**Visualization**: Stacy Gellenoncourt, Lisa A. Chakrabarti

**Writing - Original draft preparation**: Stacy Gellenoncourt, Lisa A. Chakrabarti

**Writing - Review & editing**: Stacy Gellenoncourt, Nell Saunders, Rémy Robinot, Lucas Auguste, Maaran Rajah, Jérôme Kervevan, Raphaël Jeger-Madiot, Isabelle Staropoli, Cyril Planchais, Hugo Mouquet, Julian Buchrieser, Olivier Schwartz, Lisa A. Chakrabarti

## SUPPORTING INFORMATION

**S1 Fig.**
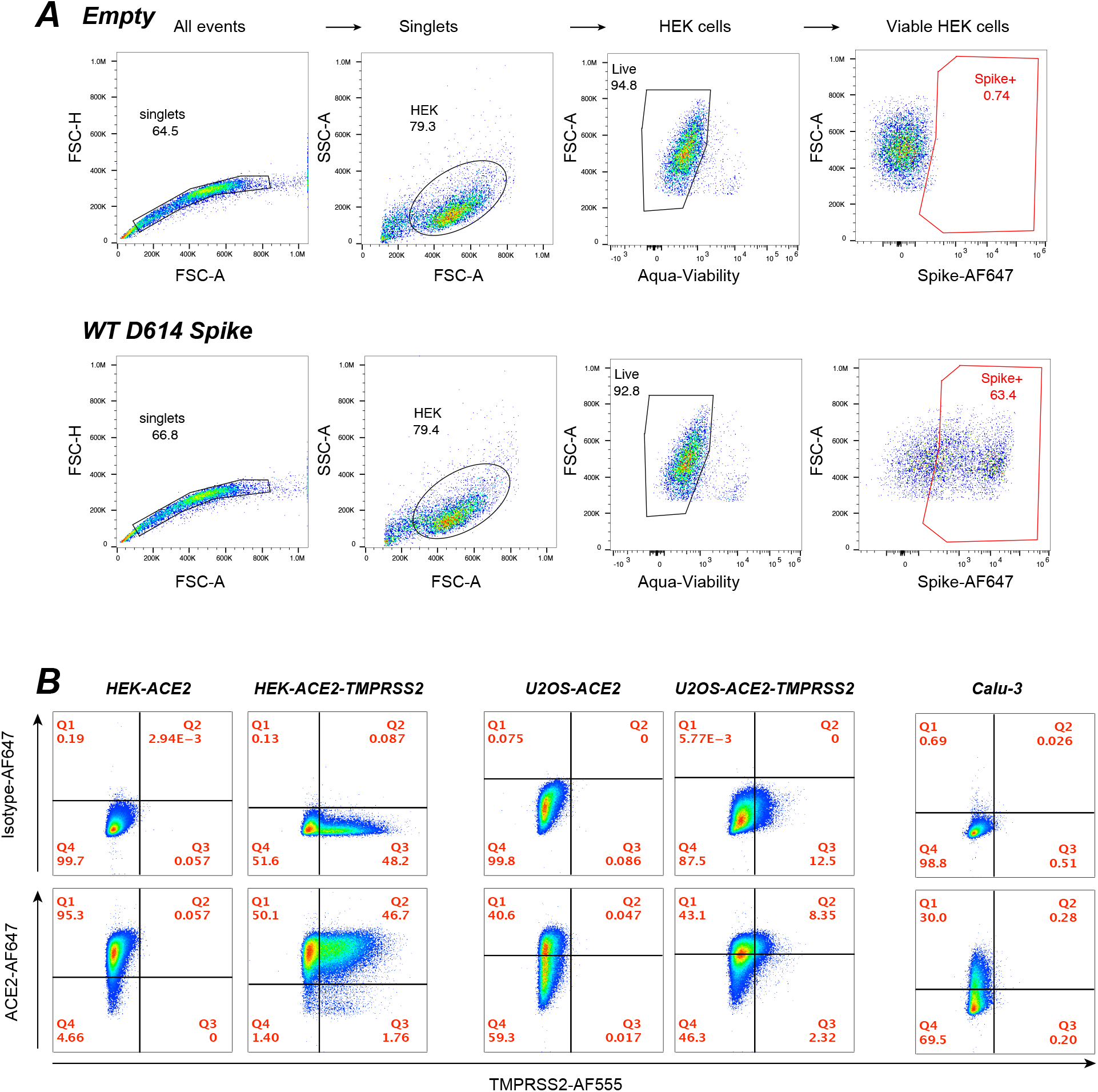
Gating strategy for spike expression and characterization of cell lines used. (A) Gating strategy to measure spike expression at the surface of transfected HEK 293Tn cells. Spike was detected in singlet viable cells after surface labeling with the human mAb 129. The top row shows cells transfected with the empty control vector, while the bottom row shows cells transfected with the WT wuhan spike vector. The percentage of spike+ cells is reported in red. (B) Expression of the receptors ACE2 and TMPRSS2 in the cell lines used in the study. Cells were surface - labeled for ACE2, permeabilized, and then labelled intracellularly for TMPRSS2. The top row shows labeling with the isotype-AF647 control, while the bottom row shows labeling with the ACE2-AF647 antibody.

**S2 Fig.**
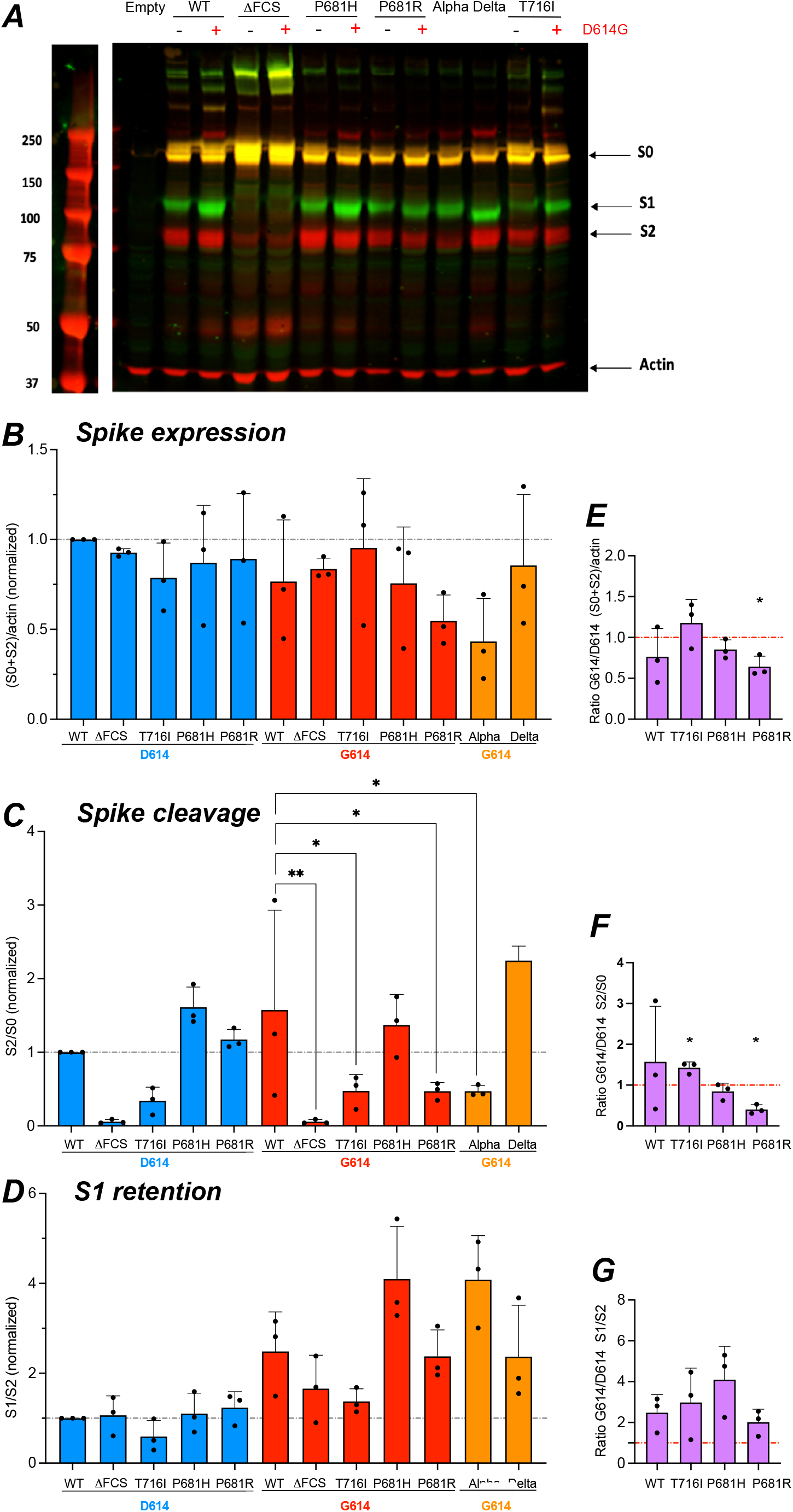
The D614 mutation promotes S1 subunit retention in producer cells. The different forms of spike present into HEK 293Tn cells transfected for GFP-pseudotyped viruses (PV) production were analyzed by Western blotting on cell lyzates. (A) Representative Western blot showing the S1 (green) and S2 (red) spike subunits, and the actin protein (red) used for normalization at the bottom of the gel. The expected size of molecular weight markers in indicated in kD on the left. The uncleaved spike precursor S0 is visible in yellow due to the superposition of green and red fluorescence. (B) Quantitation of total spike expressed (S0 + S2), reported to actin content, and normalized to WT. The fluorescence intensity of each band was quantified with the Image Studio Lite software. (C) Quantitation of total cleaved spike, measured by the S2/S0 ratio, and normalized to WT. (D) Quantitation of S1 subunit retention in producer cells, measured by the S1/S2 ratio, and normalized to WT. (B-D) Statistics were measured by one-way ANOVA, with the Holm-Sidak’s correction for multiple comparisons. Mutants in the D614 backbone were compared to the WT D614 spike, while mutants in the G614 backbone were compared to the WT G614 spike. (E-G) The ratio of total spike expressed (E), total cleaved spike (F), and S1 retention (G) measured for spikes in the G614 backbone to that in the D614 backbone is reported. Statistics evaluating whether each ratio is different from 1 were measured by one sample t-tests. (B-G) Means +/− SD are shown for n=3 independent experiments. * p<0.05; ** p<0.01.

**S3 Fig.**
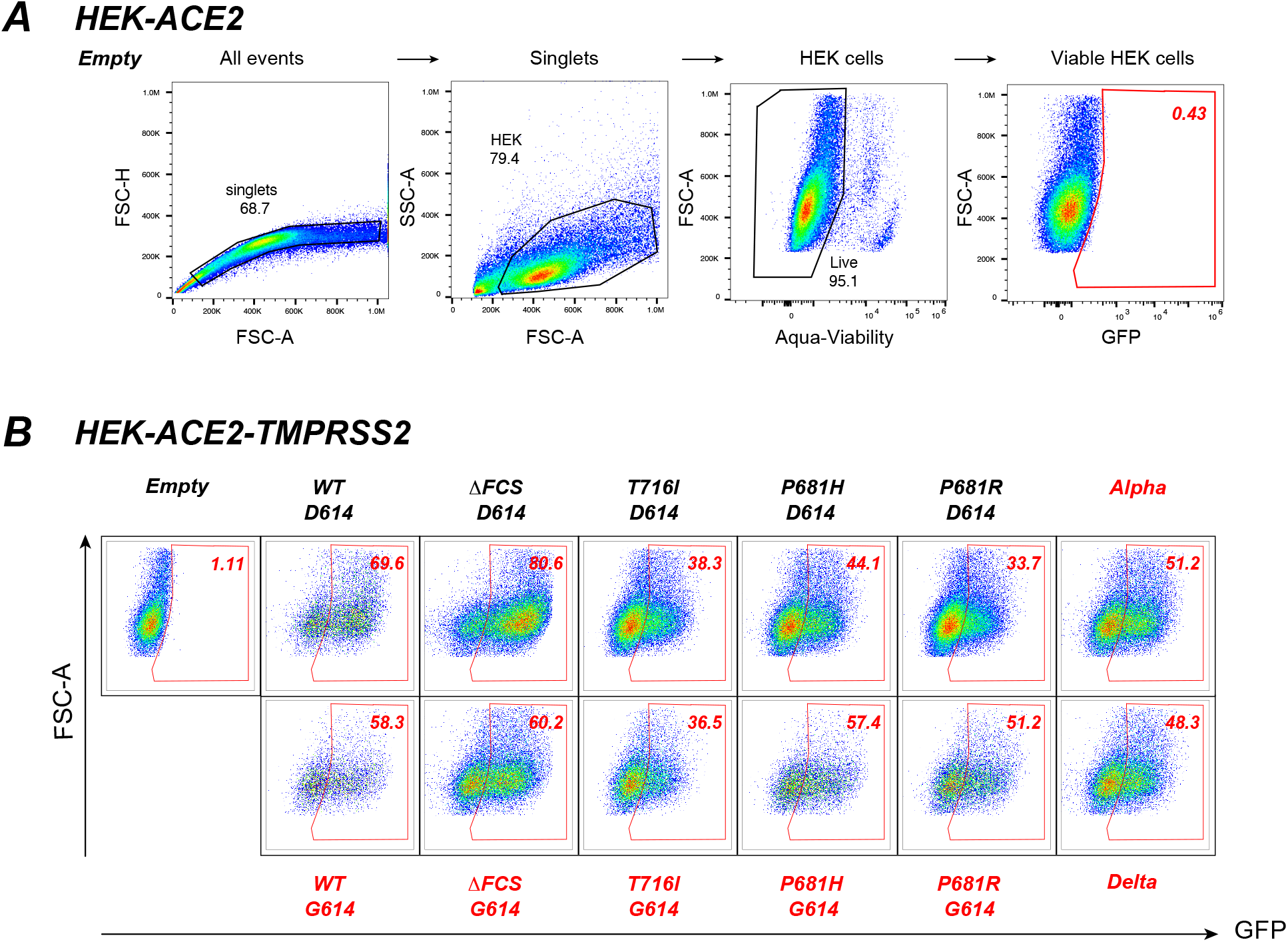
Gating strategy for the analysis of GFP-pseudovirus infectivity. (A) Gating strategy to measure the infectivity pseudotyped virus (PV) expressing the GFP reporter gene in HEK-ACE2 cells. GFP expression was evaluated at day 2 post-infection, by measuring the percentage of GFP+ cells in the singlet viable cells. A negative control corresponding to infection with 0.25 μg p24 equivalent of Empty PV is reported. (B) Infectivity of PV pseudotyped with the different spikes studied. A representative infection experiment in HEK-ACE2-TMPRSS2 cells inoculated with 0.25 μg of p24 equivalent is shown. PV infectivity is measured by the percentage of GFP+ cells reported in the top right corner of each plot.

**S4 Fig.**
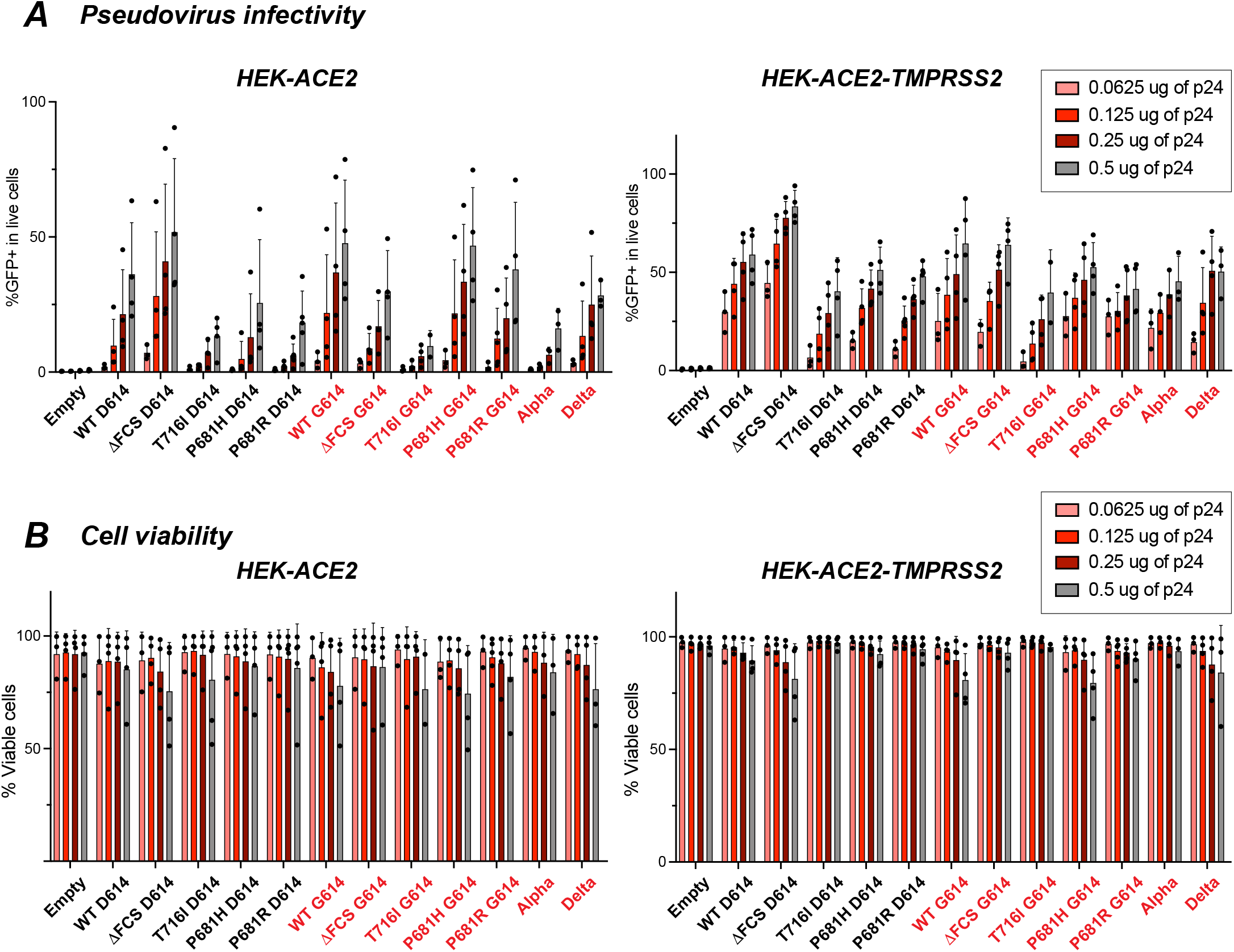
Dose response analysis of pseudovirus infectivity and target cell viability. (A) The infectivity of pseudotyped viruses (PV) carrying the different spikes was analyzed in HEK-ACE2 cells induced or not for TMPRSS2 expression. Infections were carried out with serial 2x dilutions of PV ranging from 0.5 to 0.625 μg of p24 equivalent. The percentage of GFP+ cells obtained in HEK-ACE2 cells (left) and HEK-ACE2-TMPRSS2 cells (right) is reported. (B) The percentage of viable cells, measured by exclusion of the Aqua viability dye in the singlet cell gate, is reported for each infection condition. (A,B) PV carrying a spike with the D614G mutation are labeled in red. Means +/− SD are shown for n≥3 independent experiments, except for the T716I mutant at the highest dose where n=2.

**S5 Fig.**
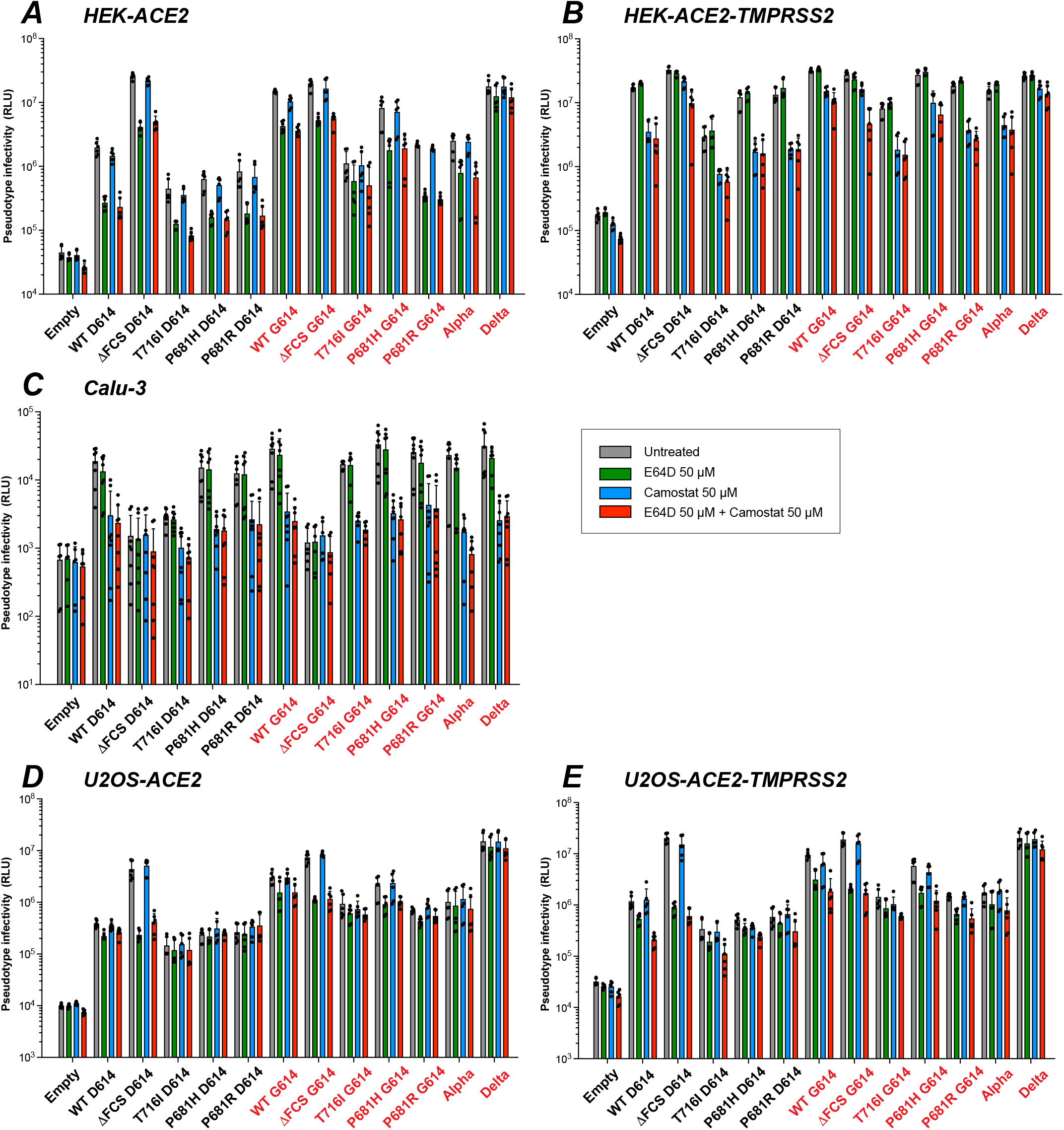
Effect of protease inhibitors on pseudovirus infectivity. The TMPRSS2 inhibitor Camostat and the cathepsin inhibitor E64D evaluated for their capacity to block pseudovtyped virus (PV) infectivity, alone or in combination. Infectivity was measured in relative luciferase units (RLU) for n=3 independent experiments, with 2 technical replicates per experiment. Conditions included untreated samples (grey) and samples treated with Camostat (blue), E64D (green), or the combination of the two inhibitors (red). Infectivity was analyzed in the following cell lines: HEK-ACE2 (A), HEK-ACE2-TMPRSS2 (B), Calu-3 (C), U2OS-ACE2 (D), and U2OS-ACE2-TMPRSS2 (E). The names of PV carrying the D614G mutation are reported in red.

**Supplementary Table 1:**
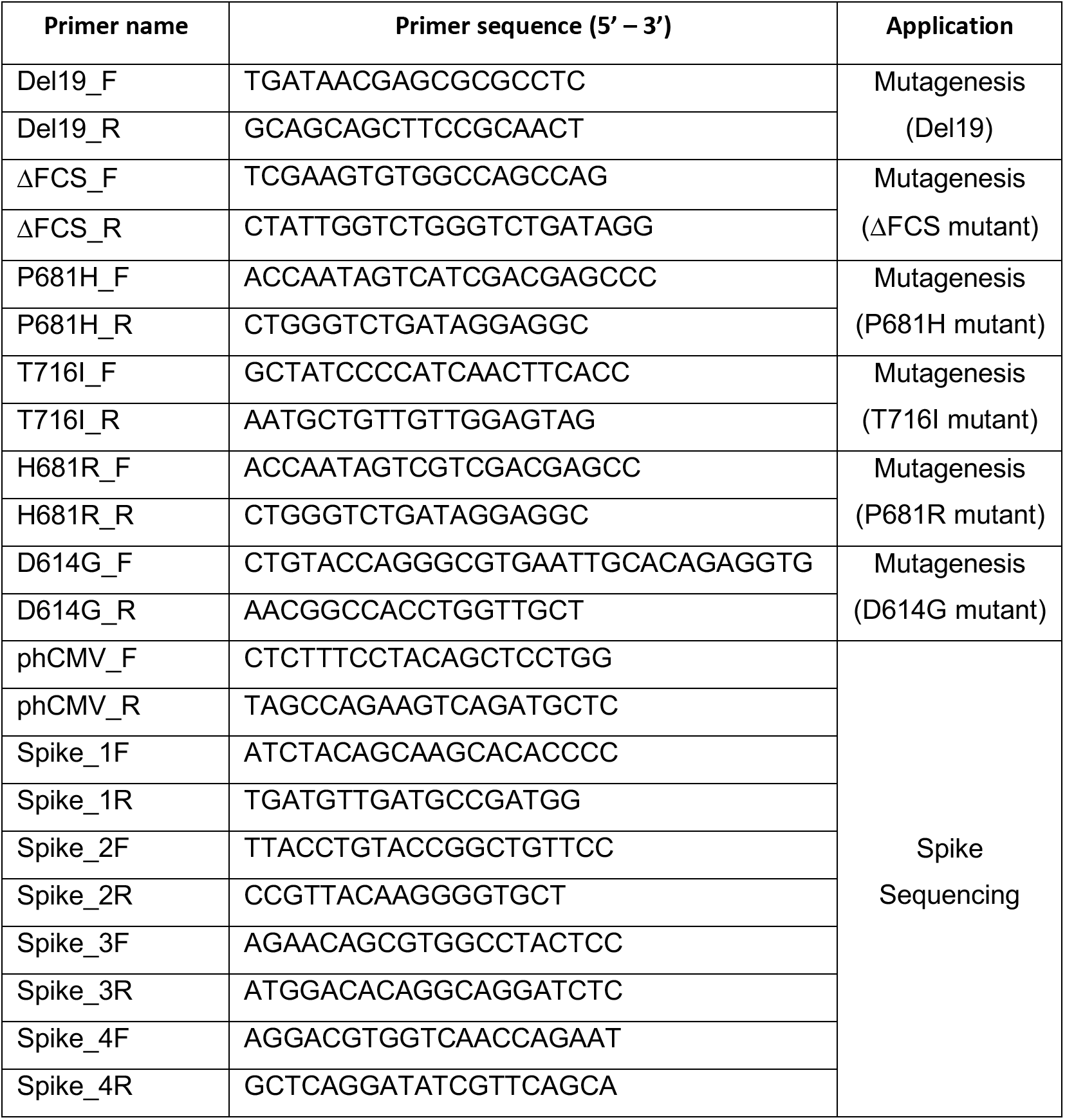
Primers used for spike mutagenesis and sequencing

## Notes

### Competing Interest Statement

The authors have declared no competing interest.

## REFERENCES

1. Tao K, Tzou PL, Nouhin J, Gupta RK, de Oliveira T, Kosakovsky Pond SL, et al. The biological and clinical significance of emerging SARS-CoV-2 variants. Nat Rev Genet. 2021;22(12):757–73. doi: 10.1038/s41576-021-00408-x. PubMed PMID: 34535792.

2. Minskaia E, Hertzig T, Gorbalenya AE, Campanacci V, Cambillau C, Canard B, et al. Discovery of an RNA virus 3’->5’ exoribonuclease that is critically involved in coronavirus RNA synthesis. Proc Natl Acad Sci U S A. 2006;103(13):5108–13. doi: 10.1073/pnas.0508200103. PubMed PMID: 16549795.

3. Korber B, Fischer WM, Gnanakaran S, Yoon H, Theiler J, Abfalterer W, et al. Tracking Changes in SARS-CoV-2 Spike: Evidence that D614G Increases Infectivity of the COVID-19 Virus. Cell. 2020. doi: 10.1016/j.cell.2020.06.043. PubMed PMID: 32697968.

4. Jackson CB, Farzan M, Chen B, Choe H. Mechanisms of SARS-CoV-2 entry into cells. Nat Rev Mol Cell Biol. 2022;23(1):3–20. doi: 10.1038/s41580-021-00418-x. PubMed PMID: 34611326.

5. WHO. Tracking SARS-CoV-2 variants 2022. Available from: https://www.who.int/en/activities/tracking-SARS-CoV-2-variants/.

6. Davies NG, Abbott S, Barnard RC, Jarvis CI, Kucharski AJ, Munday JD, et al. Estimated transmissibility and impact of SARS-CoV-2 lineage B.1.1.7 in England. Science. 2021;372(6538). doi: 10.1126/science.abg3055. PubMed PMID: 33658326.

7. Thorne LG, Bouhaddou M, Reuschl AK, Zuliani-Alvarez L, Polacco B, Pelin A, et al. Evolution of enhanced innate immune evasion by SARS-CoV-2. Nature. 2022;602(7897):487–95. doi: 10.1038/s41586-021-04352-y. PubMed PMID: 34942634.

8. Ulrich L, Halwe NJ, Taddeo A, Ebert N, Schon J, Devisme C, et al. Enhanced fitness of SARS-CoV-2 variant of concern Alpha but not Beta. Nature. 2022;602(7896):307–13. doi: 10.1038/s41586-021-04342-0. PubMed PMID: 34937050.

9. Liu L, Iketani S, Guo Y, Chan JF, Wang M, Liu L, et al. Striking Antibody Evasion Manifested by the Omicron Variant of SARS-CoV-2. Nature. 2021. doi: 10.1038/s41586-021-04388-0. PubMed PMID: 35016198.

10. Planas D, Bruel T, Grzelak L, Guivel-Benhassine F, Staropoli I, Porrot F, et al. Sensitivity of infectious SARS-CoV-2 B.1.1.7 and B.1.351 variants to neutralizing antibodies. Nat Med. 2021;27(5):917–24. doi: 10.1038/s41591-021-01318-5. PubMed PMID: 33772244.

11. Mlcochova P, Kemp S, Dhar MS, Papa G, Meng B, Ferreira I, et al. SARS-CoV-2 B.1.617.2 Delta variant replication and immune evasion. Nature. 2021. doi: 10.1038/s41586-021-03944-y. PubMed PMID: 34488225.

12. Twohig KA, Nyberg T, Zaidi A, Thelwall S, Sinnathamby MA, Aliabadi S, et al. Hospital admission and emergency care attendance risk for SARS-CoV-2 delta (B.1.617.2) compared with alpha (B.1.1.7) variants of concern: a cohort study. Lancet Infect Dis. 2022;22(1):35–42. doi: 10.1016/s1473-3099(21)00475-8. PubMed PMID: 34461056.

13. Viana R, Moyo S, Amoako DG, Tegally H, Scheepers C, Althaus CL, et al. Rapid epidemic expansion of the SARS-CoV-2 Omicron variant in southern Africa. Nature. 2022. doi: 10.1038/s41586-022-04411-y. PubMed PMID: 35042229.

14. Suzuki R, Yamasoba D, Kimura I, Wang L, Kishimoto M, Ito J, et al. Attenuated fusogenicity and pathogenicity of SARS-CoV-2 Omicron variant. Nature. 2022. doi: 10.1038/s41586-022-04462-1. PubMed PMID: 35104835.

15. Planas D, Saunders N, Maes P, Guivel-Benhassine F, Planchais C, Buchrieser J, et al. Considerable escape of SARS-CoV-2 Omicron to antibody neutralization. Nature. 2021. doi: 10.1038/s41586-021-04389-z. PubMed PMID: 35016199.

16. Wolter N, Jassat W, Walaza S, Welch R, Moultrie H, Groome M, et al. Early assessment of the clinical severity of the SARS-CoV-2 omicron variant in South Africa: a data linkage study. Lancet. 2022;399(10323):437–46. doi: 10.1016/s0140-6736(22)00017-4. PubMed PMID: 35065011.

17. Zhang L, Jackson CB, Mou H, Ojha A, Peng H, Quinlan BD, et al. SARS-CoV-2 spike-protein D614G mutation increases virion spike density and infectivity. Nat Commun. 2020;11(1):6013. doi: 10.1038/s41467-020-19808-4. PubMed PMID: 33243994.

18. Plante JA, Liu Y, Liu J, Xia H, Johnson BA, Lokugamage KG, et al. Spike mutation D614G alters SARS-CoV-2 fitness. Nature. 2020. doi: 10.1038/s41586-020-2895-3. PubMed PMID: 33106671.

19. Zhou B, Thao TTN, Hoffmann D, Taddeo A, Ebert N, Labroussaa F, et al. SARS-CoV-2 spike D614G change enhances replication and transmission. Nature. 2021;592(7852):122–7. doi: 10.1038/s41586-021-03361-1. PubMed PMID: 33636719.

20. Yurkovetskiy L, Wang X, Pascal KE, Tomkins-Tinch C, Nyalile TP, Wang Y, et al. Structural and Functional Analysis of the D614G SARS-CoV-2 Spike Protein Variant. Cell. 2020. doi: 10.1016/j.cell.2020.09.032. PubMed PMID: 32991842.

21. Benton DJ, Wrobel AG, Roustan C, Borg A, Xu P, Martin SR, et al. The effect of the D614G substitution on the structure of the spike glycoprotein of SARS-CoV-2. Proc Natl Acad Sci U S A. 2021;118(9). doi: 10.1073/pnas.2022586118. PubMed PMID: 33579792.

22. Zhang J, Cai Y, Xiao T, Lu J, Peng H, Sterling SM, et al. Structural impact on SARS-CoV-2 spike protein by D614G substitution. Science. 2021;372(6541):525–30. doi: 10.1126/science.abf2303. PubMed PMID: 33727252.

23. Liu Y, Liu J, Plante KS, Plante JA, Xie X, Zhang X, et al. The N501Y spike substitution enhances SARS-CoV-2 infection and transmission. Nature. 2022;602(7896):294–9. doi: 10.1038/s41586-021-04245-0. PubMed PMID: 34818667.

24. Coutard B, Valle C, de Lamballerie X, Canard B, Seidah NG, Decroly E. The spike glycoprotein of the new coronavirus 2019-nCoV contains a furin-like cleavage site absent in CoV of the same clade. Antiviral Res. 2020. doi: 10.1016/j.antiviral.2020.104742. PubMed PMID: 32057769.

25. Papa G, Mallery DL, Albecka A, Welch LG, Cattin-Ortola J, Luptak J, et al. Furin cleavage of SARS-CoV-2 Spike promotes but is not essential for infection and cell-cell fusion. PLoS Pathog. 2021;17(1):e1009246. doi: 10.1371/journal.ppat.1009246. PubMed PMID: 33493182.

26. Shang J, Wan Y, Luo C, Ye G, Geng Q, Auerbach A, et al. Cell entry mechanisms of SARS-CoV-2. Proc Natl Acad Sci U S A. 2020;117(21):11727–34. doi: 10.1073/pnas.2003138117. PubMed PMID: 32376634.

27. Hoffmann M, Kleine-Weber H, Pohlmann S. A Multibasic Cleavage Site in the Spike Protein of SARS-CoV-2 Is Essential for Infection of Human Lung Cells. Mol Cell. 2020. doi: 10.1016/j.molcel.2020.04.022. PubMed PMID: 32362314.

28. Peacock TP, Goldhill DH, Zhou J, Baillon L, Frise R, Swann OC, et al. The furin cleavage site in the SARS-CoV-2 spike protein is required for transmission in ferrets. Nat Microbiol. 2021. doi: 10.1038/s41564-021-00908-w. PubMed PMID: 33907312.

29. Johnson BA, Xie X, Bailey AL, Kalveram B, Lokugamage KG, Muruato A, et al. Loss of furin cleavage site attenuates SARS-CoV-2 pathogenesis. Nature. 2021;591(7849):293–9. doi: 10.1038/s41586-021-03237-4. PubMed PMID: 33494095.

30. Benton DJ, Wrobel AG, Xu P, Roustan C, Martin SR, Rosenthal PB, et al. Receptor binding and priming of the spike protein of SARS-CoV-2 for membrane fusion. Nature. 2020. doi: 10.1038/s41586-020-2772-0. PubMed PMID: 32942285.

31. Hoffmann M, Kleine-Weber H, Schroeder S, Kruger N, Herrler T, Erichsen S, et al. SARS-CoV-2 Cell Entry Depends on ACE2 and TMPRSS2 and Is Blocked by a Clinically Proven Protease Inhibitor. Cell. 2020. doi: 10.1016/j.cell.2020.02.052. PubMed PMID: 32142651.

32. Koch J, Uckeley ZM, Doldan P, Stanifer M, Boulant S, Lozach PY. TMPRSS2 expression dictates the entry route used by SARS-CoV-2 to infect host cells. EMBO J. 2021;40(16):e107821. doi: 10.15252/embj.2021107821. PubMed PMID: 34159616.

33. Tian S, Huang Q, Fang Y, Wu J. FurinDB: A database of 20-residue furin cleavage site motifs, substrates and their associated drugs. Int J Mol Sci. 2011;12(2):1060–5. doi: 10.3390/ijms12021060. PubMed PMID: 21541042.

34. Brown JC, Goldhill DH, Zhou J, Peacock TP, Frise R, Goonawardane N, et al. Increased transmission of SARS-CoV-2 lineage B.1.1.7 (VOC 2020212/01) is not accounted for by a replicative advantage in primary airway cells or antibody escape. bioRxiv. 2021:2021.02.24.432576. doi: 10.1101/2021.02.24.432576.

35. Escalera A, Gonzalez-Reiche AS, Aslam S, Mena I, Laporte M, Pearl RL, et al. Mutations in SARS-CoV-2 variants of concern link to increased spike cleavage and virus transmission. Cell Host Microbe. 2022. doi: 10.1016/j.chom.2022.01.006. PubMed PMID: 35150638.

36. Rajah MM, Hubert M, Bishop E, Saunders N, Robinot R, Grzelak L, et al. SARS-CoV-2 Alpha, Beta, and Delta variants display enhanced Spike-mediated syncytia formation. EMBO J. 2021;40(24):e108944. doi: 10.15252/embj.2021108944. PubMed PMID: 34601723.

37. Lubinski B, Fernandes MHV, Frazier L, Tang T, Daniel S, Diel DG, et al. Functional evaluation of the P681H mutation on the proteolytic activation of the SARS-CoV-2 variant B.1.1.7 (Alpha) spike. iScience. 2022;25(1):103589. doi: 10.1016/j.isci.2021.103589. PubMed PMID: 34909610.

38. Liu Y, Liu J, Johnson BA, Xia H, Ku Z, Schindewolf C, et al. Delta spike P681R mutation enhances SARS-CoV-2 fitness over Alpha variant. bioRxiv. 2021:2021.08.12.456173. doi: 10.1101/2021.08.12.456173.

39. Peacock TP, Sheppard CM, Brown JC, Goonawardane N, Zhou J, Whiteley M, et al. The SARS-CoV-2 variants associated with infections in India, B.1.617, show enhanced spike cleavage by furin. bioRxiv. 2021:2021.05.28.446163. doi: 10.1101/2021.05.28.446163.

40. Saito A, Irie T, Suzuki R, Maemura T, Nasser H, Uriu K, et al. Enhanced fusogenicity and pathogenicity of SARS-CoV-2 Delta P681R mutation. Nature. 2022;602(7896):300–6. doi: 10.1038/s41586-021-04266-9. PubMed PMID: 34823256.

41. Meng B, Abdullahi A, Ferreira I, Goonawardane N, Saito A, Kimura I, et al. Altered TMPRSS2 usage by SARS-CoV-2 Omicron impacts tropism and fusogenicity. Nature. 2022. doi: 10.1038/s41586-022-04474-x. PubMed PMID: 35104837.

42. Hui KPY, Ho JCW, Cheung MC, Ng KC, Ching RHH, Lai KL, et al. SARS-CoV-2 Omicron variant replication in human bronchus and lung ex vivo. Nature. 2022. doi: 10.1038/s41586-022-04479-6. PubMed PMID: 35104836.

43. Rajah MM, Bernier A, Buchrieser J, Schwartz O. The Mechanism and Consequences of SARS-CoV-2 Spike-Mediated Fusion and Syncytia Formation. J Mol Biol. 2021:167280. doi: 10.1016/j.jmb.2021.167280. PubMed PMID: 34606831.

44. Klein S, Cortese M, Winter SL, Wachsmuth-Melm M, Neufeldt CJ, Cerikan B, et al. SARS-CoV-2 structure and replication characterized by in situ cryo-electron tomography. Nat Commun. 2020;11(1):5885. doi: 10.1038/s41467-020-19619-7. PubMed PMID: 33208793.

45. Nguyen HT, Zhang S, Wang Q, Anang S, Wang J, Ding H, et al. Spike glycoprotein and host cell determinants of SARS-CoV-2 entry and cytopathic effects. J Virol. 2020. doi: 10.1128/JVI.02304-20. PubMed PMID: 33310888.

46. Gu Y, Cao J, Zhang X, Gao H, Wang Y, Wang J, et al. Receptome profiling identifies KREMEN1 and ASGR1 as alternative functional receptors of SARS-CoV-2. Cell Res. 2022;32(1):24–37. doi: 10.1038/s41422-021-00595-6. PubMed PMID: 34837059.

47. Gobeil SM, Janowska K, McDowell S, Mansouri K, Parks R, Stalls V, et al. Effect of natural mutations of SARS-CoV-2 on spike structure, conformation, and antigenicity. Science. 2021;373(6555). doi: 10.1126/science.abi6226. PubMed PMID: 34168071.

48. Meng B, Kemp SA, Papa G, Datir R, Ferreira I, Marelli S, et al. Recurrent emergence of SARS-CoV-2 spike deletion H69/V70 and its role in the Alpha variant B.1.1.7. Cell Rep. 2021;35(13):109292. doi: 10.1016/j.celrep.2021.109292. PubMed PMID: 34166617.

49. Bussani R, Schneider E, Zentilin L, Collesi C, Ali H, Braga L, et al. Persistence of viral RNA, pneumocyte syncytia and thrombosis are hallmarks of advanced COVID-19 pathology. EBioMedicine. 2020;61:103104. doi: 10.1016/j.ebiom.2020.103104. PubMed PMID: 33158808.

50. Davies NG, Jarvis CI, Group CC-W, Edmunds WJ, Jewell NP, Diaz-Ordaz K, et al. Increased mortality in community-tested cases of SARS-CoV-2 lineage B.1.1.7. Nature. 2021;593(7858):270–4. doi: 10.1038/s41586-021-03426-1. PubMed PMID: 33723411.

51. McCallum M, Czudnochowski N, Rosen LE, Zepeda SK, Bowen JE, Walls AC, et al. Structural basis of SARS-CoV-2 Omicron immune evasion and receptor engagement. Science. 2022;375(6583):864–8. doi: 10.1126/science.abn8652. PubMed PMID: 35076256.

52. Zhang J, Cai Y, Lavine CL, Peng H, Zhu H, Anand K, et al. Structural and functional impact by SARS-CoV-2 Omicron spike mutations. bioRxiv. 2022:2022.01.11.475922. doi: 10.1101/2022.01.11.475922.

53. Boson B, Legros V, Zhou B, Siret E, Mathieu C, Cosset FL, et al. The SARS-CoV-2 envelope and membrane proteins modulate maturation and retention of the spike protein, allowing assembly of virus-like particles. J Biol Chem. 2021;296:100111. doi: 10.1074/jbc.RA120.016175. PubMed PMID: 33229438.

54. Rocheleau L, Laroche G, Fu K, Stewart CM, Mohamud AO, Côté M, et al. Identification of a High-Frequency Intrahost SARS-CoV-2 Spike Variant with Enhanced Cytopathic and Fusogenic Effects. mBio. 2021;12(3):e0078821. doi: 10.1128/mBio.00788-21. PubMed PMID: 34182784.

55. Buchrieser J, Dufloo J, Hubert M, Monel B, Planas D, Rajah MM, et al. Syncytia formation by SARS-CoV-2-infected cells. EMBO J. 2021;40(3):e107405. doi: 10.15252/embj.2020107405. PubMed PMID: 33522642.

56. Crawford KHD, Eguia R, Dingens AS, Loes AN, Malone KD, Wolf CR, et al. Protocol and Reagents for Pseudotyping Lentiviral Particles with SARS-CoV-2 Spike Protein for Neutralization Assays. Viruses. 2020;12(5). doi: 10.3390/v12050513. PubMed PMID: 32384820.

